# Structural assemblies for an RNA world

**DOI:** 10.64898/2026.07.01.735769

**Authors:** Yangyi Ren, Zhenglai Zhang, Ke Chen, Mengxiao Li, Yuting Xie, Tiantian Bai, Baowei Huang, Bowen Xiao, Eric Westhof, David M J Lilley, Jia Wang, Zhichao Miao, Xuepeng Wei, Lin Huang

**Author notes:** These authors contributed equally (Yangyi Ren, Zhenglai Zhang, Ke Chen).

## Abstract

Protein homo-oligomerization widely generates symmetric assemblies for encapsulation and scaffolding. Conversely, natural RNA quaternary structures appear limited, with known RNA-only multimers predominantly forming dimers or dihedral assemblies from long transcripts. Whether short RNAs can access broader higher-order assembly space has remained unclear. Here, using cryo-EM, we show that RNAs under 200 nucleotides form three distinct structural classes: a 60-subunit viral-capsid-like icosahedron, non-caged oligomers including a trimer and a strand-exchanged dimer, and a continuous filament assembled from a 57-nucleotide RNA. These structures demonstrate that assembly complexity does not simply scale with RNA length; compact RNAs can specify architectures traditionally associated with proteins. Our findings broaden the known RNA quaternary repertoire, strengthen the structural plausibility of higher-order organization in an RNA world, and establish foundational reference architectures to guide future RNA-based self-assembling nanoparticle design.

**Highlights:**

- Short RNAs reveal protein-like assembly principles for an RNA world
- The first natural RNA-only T=1 icosahedral cage is a viral-capsid-like 60-mer
- Compact RNA–RNA interfaces encode global curvature, valency and propagation
- RNA motifs assemble into closed cages, finite oligomers and extended filaments

## INTRODUCTION

Protein homo-oligomerization is a pervasive organizing principle in biology.^1–4^ By assembling identical subunits into rings, cages, tubes, and filaments, proteins generate architectures that support encapsulation, scaffolding, multivalency, transport of cargo, and long-range cooperativity.^5–9^ Such assemblies underlie functions as diverse as cytoskeletal organization and polyhedral membrane coats, and across proteomes, they represent a foundational route to biological structure and function.^8,10,11^ If early life passed through an RNA world, however, comparable forms of higher-order organization would have had to emerge without proteins.^12,13^ This raises a fundamental question: can RNA access broad assembly regimes as seen in protein biology?

RNA, by contrast, has long been viewed primarily through the lens of tertiary folding rather than quaternary assembly.^14,15^ Although RNA can form intricate three-dimensional structures and engage in kissing-loop and pseudoknot formation, co-axial stacking, and many other tertiary contacts, naturally occurring higher-order RNA assemblies are limited compared with proteins.^16–18^ Reviews of RNA quaternary structure have therefore emphasized a striking difference between the protein and RNA worlds: proteins routinely adopt closed and open assembly geometries, whereas natural RNAs seem largely confined to monomers, dimers, or a few specialized oligomers.^19–23^

Recent cryo-EM studies have begun to lead to a revision of this view. Structures of OLE, ROOL, GOLLD, ARRPOF, and related RNAs have shown that natural pure RNA (i.e. free from protein) can form symmetric multimers and nanocage-like assemblies^24–29^(Figure 1A, left). Yet these examples still occupy a relatively narrow region of assembly space. OLE and ARRPOF remain dimeric, whereas ROOL and GOLLD assemble into dihedral multimers built from half-shell layers coupled by interplanar contacts. Because these systems are also predominantly assembled from longer RNA protomers, typically a few hundred nucleotides, it has been assumed that multimeric assembly requires longer RNA molecules (Figure 1B, green circles).

**Figure 1.**
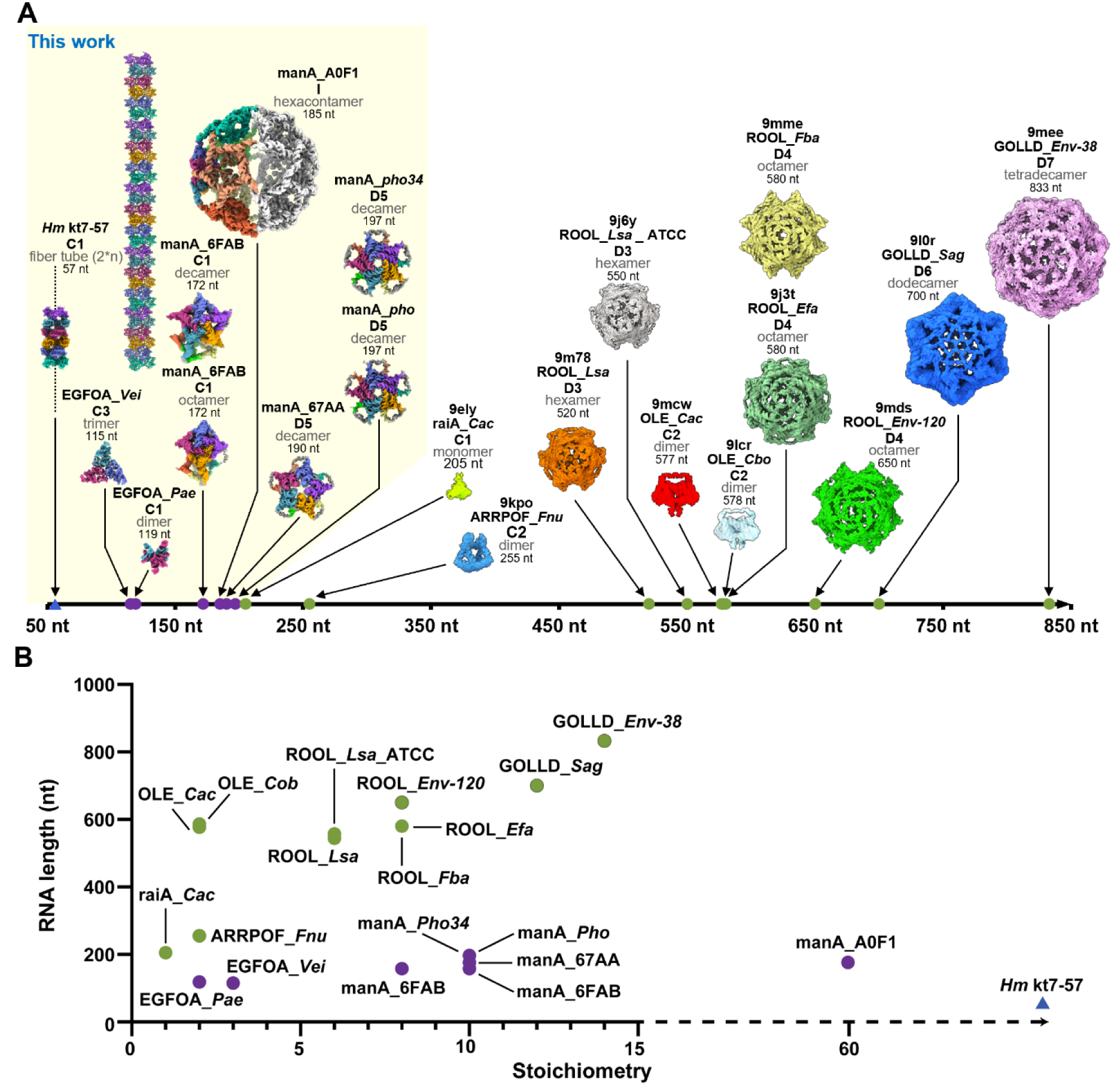
Short RNAs access major architectural regimes of higher-order assembly. (**A**) Gallery of self-assembling RNA structures comparing those determined in this study (left, highlighted in light yellow) with previously reported natural RNA assemblies (right). Structures are arranged along a timeline according to their monomeric nucleotide length (nt). (**B**) Scatter plot correlating RNA monomer length with assembly stoichiometry. Previous structures (green circles) typically show that higher oligomerization requires longer RNA sequences, whereas the assemblies reported here are highlighted in purple circles and the blue triangle.

A second unresolved issue concerns the architectural level. Lower-order symmetric assemblies frequently occur in cyclic or dihedral forms,^30^ whereas a 60-subunit closed shell belongs to a more demanding assembly class that must satisfy global curvature and closure under icosahedral constraints^31^ (Figure S1). The distinction is therefore not simply quantitative. Whether even long RNAs can access this level of organization has remained uncertain, let alone short ones.

This question bears directly on the plausibility of the RNA world. RNA-world models require RNA not only to store information and catalyze reactions but also to generate higher-order organization.^32,33^ Cech argued that some form of encapsulation was likely a key early step in the origins of life because it can protect genomes from degradation, compartmentalize and concentrate metabolite molecules, and allow selection to act on molecular systems as units^34^. More broadly, the plausibility of an RNA world depends on whether such functional and structural capacities can emerge from RNA sequence space (Figure S2).^35,36^

This issue has become timely. A recent study described QT45, a 45-nucleotide polymerase ribozyme that can synthesize both its complementary strand and a copy of itself, arguing that a key catalytic function required for the RNA world can reside in a surprisingly small RNA element and suggesting that such activities may be more abundant in RNA sequence space than previously thought.^35^ By analogy, if short RNAs can also encode complex functions, then another major requirement of an RNA world, quaternary assembly, may similarly be accessible to relatively small RNAs.

Here we show that this is the case. Using cryo-EM, we identify higher-order assemblies formed by RNAs shorter than 200 nucleotides that span three major structural regimes: closed shells, defined non-cage oligomers, and open-ended filaments. These architectures are specified by compact RNA-RNA interaction modules rather than by the folding capacity of long transcripts. Our findings expand the structural repertoire of RNA, show that short RNAs can form assemblies long associated with proteins, and thereby strengthen the structural plausibility of an RNA world. Beyond their evolutionary significance, they can also serve as an additional source of inspiration for the design of RNA-based self-assembling nanoparticles in biomedical applications.^19,37,38^

## RESULTS

### Short RNAs assemble into diverse architectures, from closed cages to extended filaments

To test whether short RNAs can access higher-order assembly regimes analogous to those commonly observed in proteins, we analyzed three unrelated RNA families by cryo-EM. The manA family (RF01745)^39,40^ forms discrete closed oligomers, including an unexpected 60-subunit icosahedral-like cage. Members of the EGFOA family (RF02955)^41^ populate distinct non-cage oligomeric states, including a stable trimer and a strand-exchanged dimer. The 57-nt *Hm* kt7-57 based on the kink-turn motif,^42,43^ forms a continuous RNA-only filament through repeated intermolecular interactions. Together, these structures define three distinct assembly regimes for short RNAs—closed, non-cage, and open-ended—leading to the analyses that follow.

### Compact interaction modules override RNA length in determining assembly scale

Placing these structures alongside previously characterized natural RNA oligomers reveals a shift in how RNA assembly should be understood (Figure 1 and Figure S3). Recent structures of OLE, ROOL and GOLLD established that protein-free natural RNAs can form symmetric multimers and nanocage-like assemblies. However, these examples are built from substantially longer protomers, typically several hundred nucleotides in length, and the only reported assemblies with more than two subunits have been ROOL and GOLLD. Taken together, these observations have encouraged the view that increasing assembly complexity follows increasing RNA length (Figure 1B, green circles). By contrast, the assemblies reported here are formed by RNAs shorter than 200 nt yet span three major structural regimes: a closed 60-subunit icosahedral-like shell, defined non-cage oligomers including trimers and strand-exchanged dimers, and an open-ended filament. These comparisons indicate that assembly complexity in natural RNAs is governed less by transcript length itself than by compact interaction geometries that specify curvature, valency or axial propagation.

### The manA RNA motif forms a 60-mer icosahedral-like cage and smaller closed oligomers

The manA RNA motif is a conserved putative cis-regulatory element frequently found in the 5′ untranslated regions of genes in *Photobacterium* species or cyanobacteria-infecting phages.^39^ Cryo-EM analysis showed that related manAs do not adopt a single oligomeric state, but rather assemble into multiple discrete closed architectures. The most striking example was manA_A0F1, which formed hollow spherical particles readily visible in cryo-EM micrographs and yielded a 60-subunit reconstruction with an overall diameter of ∼430 Å (Figure 2A, B, and Figure S4). The assembly adopts a truncated icosahedral geometry resembling a fullerene-like cage (Figure 2C). Structural analysis further showed that the assembly adopts a T=1 icosahedral geometry. Contextualizing this architecture alongside naturally occurring protein cages demonstrates that its scale and complexity bridge the gap between viral capsids (STNV and AAV) and D6-symmetric cellular machinery, such as the clathrin cage (Figure 2D–G and Figure S5).

**Figure 2.**
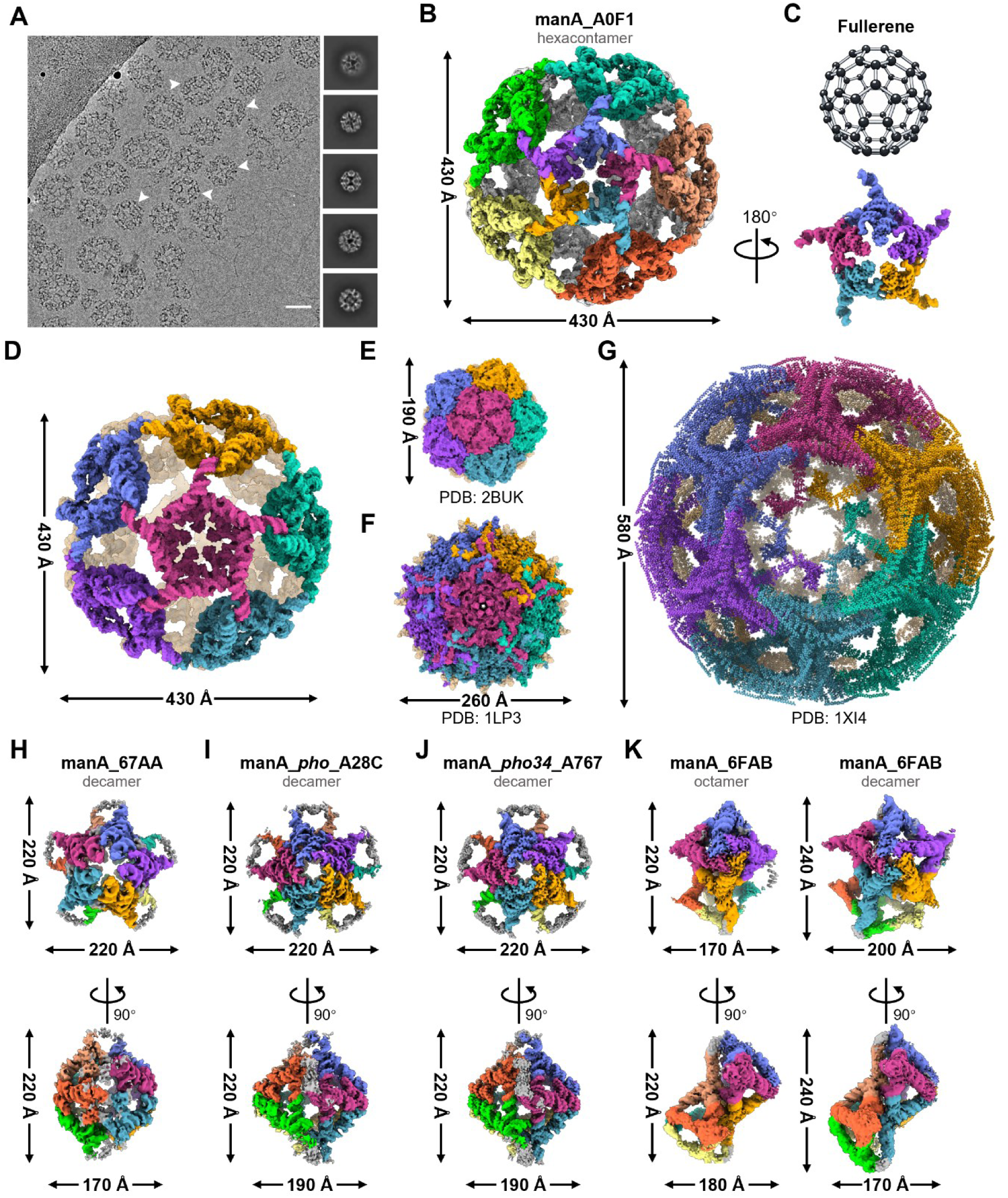
Cryo-EM structure of a fullerene-like RNA hexacontamer and its oligomeric variants. (**A**) Representative cryo-EM micrograph of the manA_A0F1. White arrowheads indicate individual spherical particles. The right panels show representative 2D class averages. Scale bar, 40 nm. Although many particles converged into a homogeneous spherical class (∼430 Å in diameter) used for high-resolution reconstruction, the dataset also contained larger and morphologically heterogeneous particles (∼400–750 Å overall), many of which appeared ellipsoidal or flattened and were excluded during classification. (**B**) Cryo-EM density map of the manA_A0F1 hexacontamer (60-mer) with an overall diameter of ∼430 Å. (**C**) Structural comparison with a C₆₀ fullerene cage (top), highlighting the conserved truncated icosahedral geometry. A constituent pentameric building block extracted from the assembly is shown below. (**D**) Surface representation of the complete manA_A0F1 60-mer assembly, exhibiting a T=1 icosahedral geometry. (**E** to **G**) Structural and dimensional comparisons with naturally occurring protein cages to provide scale and geometric context. Shown are surface models of representative T=1 icosahedral viral capsids, Satellite Tobacco Necrosis Virus (STNV, PDB: 2BUK) (**E**) and Adeno-Associated Virus (AAV, PDB: 1LP3) (**F**), alongside a larger D6-symmetric clathrin cage (PDB 1XI4) (**G**). (**H** to **K**) Top and side views (rotated by 90°) of the decameric assemblies for manA_67AA (**H**), manA_*pho*_A28C (**I**), and manA_*pho34*_A767 (**J**). (**K**) Structural polymorphism observed in manA_6FAB, which forms both octameric (left) and decameric (right) assemblies. Overall dimensions for each assembly are indicated.

Other manA variants adopted smaller but still discrete oligomeric states. manA_67AA, manA_*pho*_A28C, and manA_*pho34*_A767 formed decameric assemblies, whereas manA_6FAB exhibited structural polymorphism and adopted both octameric and decameric structures (Figure 2H-K and Figure S6-12). The coexistence of these related but distinct states indicates that the manA scaffold is intrinsically polymorphic, yet does not simply aggregate into poorly defined particles. Instead, closely related short RNAs encode specific folds with oriented interaction modules that converge on defined stoichiometries.

### Distinct intermolecular interfaces share a pentameric building block in manA assemblies

Despite their different final stoichiometries, the decameric and 60-subunit manA assemblies are built from a common pentameric building block (Figure 3A, B). On the basis of sequence and structural comparisons, we have grouped the experimentally characterized variants into two related sets that retain a conserved core fold that then differ in their peripheral helices (Figure S13-16). In manA_A0F1, the pentamer is relatively flat and is stabilized primarily by two intermolecular kissing-loop interfaces, L5-L7′ and L8-L9′ (Figure 3C-E, Figure S13 and Supplementary Figure S17A, B). These interactions organize five monomers into a star-shaped pentamer that serves as the basic module of the 60-mer cage.

**Figure 3.**
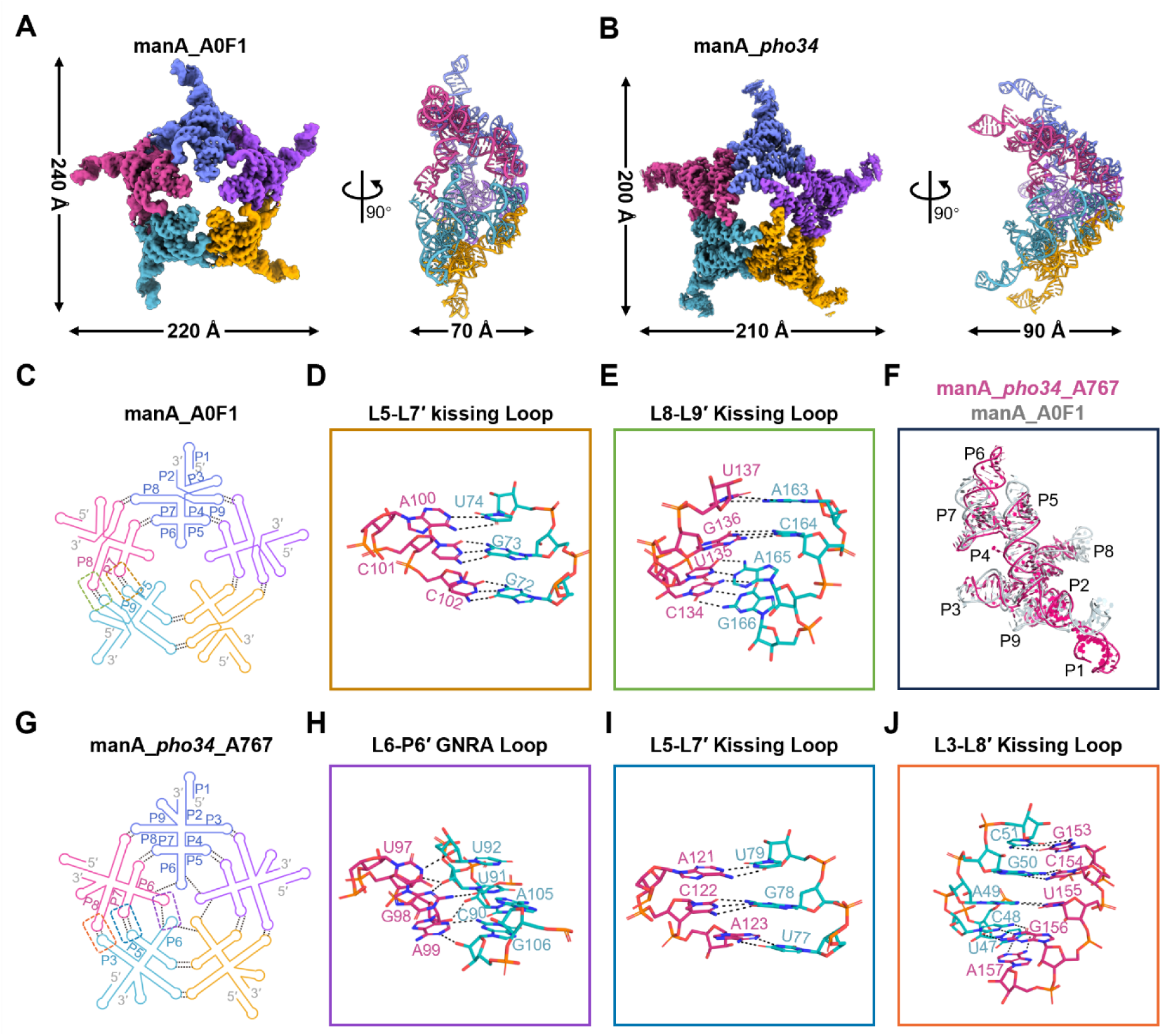
Pentameric building blocks stabilized by extensive intermolecular kissing loops and tertiary interactions. (**A** and **B**) Cryo-EM density maps and atomic models of the pentameric building blocks for manA_A0F1 (**A**) and manA_*pho34*_A767 (**B**). Top and side views (rotated by 90°) are shown, with overall dimensions indicated. (**C**) Projected secondary structure schematic of the manA_A0F1 pentamer. Colored dashed boxes indicate the interfaces detailed in (**D**) and (**E**). (**D** and **E**) Close-up atomic views of the intermolecular L5-L7′ kissing loop (**D**) and L8-L9′ kissing loop (**E**) interactions that stabilize the A0F1 pentamer. Hydrogen bonds are shown as black dashed lines. (**F**) Structural superposition of the manA_*pho34*_A767 (pink) and manA_A0F1 (gray) monomers. (**G**) Projected secondary structure schematic of the manA_*pho34*_A767 pentamer. (**H** to **J**) Detailed atomic views of the specific tertiary interactions mediating the manA_*pho34*_A767 assembly, including the L6-P6′ GNRA loop-receptor interaction (**H**), the L5-L7′ kissing loop (**I**), and the L3-L8′ kissing loop (**J**).

A related but distinct pentameric architecture is formed by manA_*pho34*_A767. Structural superposition showed that the monomeric cores of manA_A0F1 and manA_*pho34*_A767 are similar. In contrast, their peripheral helices adopt different spatial arrangements resulting in a dome-shaped pentameric assembly (Figure 3F). In manA_*pho34*_A767, pentamer formation is not driven by the same pair of interfaces used in A0F1, but rather by an alternative interaction network involving the L6-P6′ GNRA loop–receptor contact, together with L5-L7′ and L3-L8′ kissing-loop interactions (Figure 3G-J and Figure S17 11D-F). Thus, related manA RNAs converge on shared pentameric building blocks through nonidentical local base pairs. In this family, geometric reuse of a pentameric intermediate, rather than RNA length alone, appears to underlie access to both smaller closed oligomers and the large 60-subunit cage.

### EGFOA RNAs populate non-cage oligomers centered on a stable trimeric core

To determine whether short RNAs can also form higher-order assemblies beyond closed-cage geometries, we examined the EGFOA RNA family, a set of conserved, putative small RNAs found in unusually large intergenic regions lacking annotated protein-coding genes.^41^ Cryo- EM analysis of EGFOA_*Vei*_C1A6 revealed a stable trimeric assembly at 2.96 Å resolution (Figure 4A and Figure S18). The trimer adopts a bent, boomerang-like architecture rather than a closed cage, establishing a non-cage assembly regime for short natural RNAs. Its structural integrity is maintained by an extensive intermolecular network that includes Watson-Crick and non-Watson-Crick base pairing, ribose-phosphate hydrogen bonding, and base stacking (Figure 4C-F and Figure S19).

**Figure 4.**
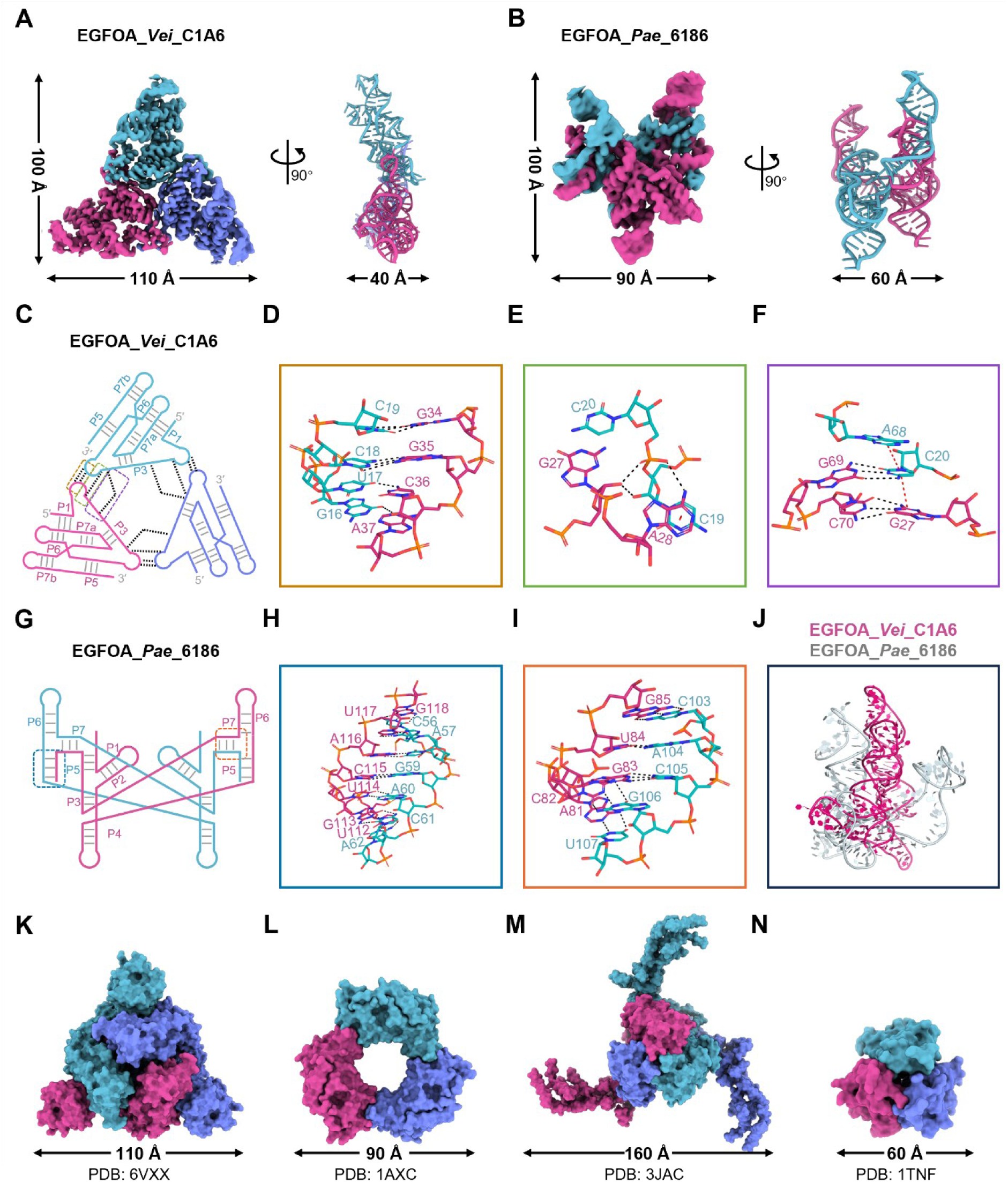
EGFOA RNAs define a non-cage oligomeric regime. (**A** and **B**) Cryo-EM density maps and the corresponding atomic models of the EGFOA_*Vei*_C1A6 trimer (**A**) and EGFOA_*Pae*_6186 dimer (**B**). Top and side views (rotated by 90°) are shown. (**C**) Projected secondary structure schematic of the EGFOA_*Vei*_C1A6 trimer, illustrating the specific intermolecular interaction network. Colored dashed boxes indicate the interfaces detailed in (**D**) to (**F)**. (**D** to **F**) Detailed atomic views of the interaction network stabilizing the trimer, including canonical and non-canonical base pairing (**D**), extensive hydrogen bonds between the ribose sugar and phosphate backbone (**E**), and complex base stacking involving π-π interactions (indicated by red dashed lines) alongside standard hydrogen bonds (black dashed lines) (**F**). (**G**) Projected secondary structure schematic of the EGFOA_*Pae*_6186 dimer, highlighting the extensive intermolecular interfaces. (**H** and **I**) Close-up atomic views of the intermolecular P5 (**H**) and P7 (**I**) helical stems, which are formed via 3′-terminal strand displacement (strand exchange) between the two interacting monomers. (**J**) Structural superposition of the constituent EGFOA_*Vei*_C1A6 (pink) and EGFOA_*Pae*_6186 (gray) monomers. (**K** to **N**) Structural comparison of the EGFOA RNA trimer with representative naturally occurring C3-symmetric proteins. Surface models (top views) are shown for the SARS-CoV-2 spike protein (PDB 6VXX) (**K**), proliferating cell nuclear antigen (PCNA, PDB 1AXC) (**I**), the entire mechanosensitive Piezo1 channel (PDB 3JAC) (**M**), and the soluble cell-signaling cytokine tumor necrosis factor-α (TNF-α, PDB 1TNF) (**N**).

Three-dimensional classification identified an additional oligomeric state, a lower-resolution hexamer, indicating that the trimer can serve as a building block for larger assemblies (Figure S20). In contrast to the bounded icosahedral-like shell formed by manA_A0F1, the EGFOA_*Vei*_C1A6 trimer shows that short RNAs can also adopt non-cage topologies that more closely resemble the diverse planar or open oligomeric arrangements commonly found in protein homo-oligomers (Figure 4K-N).^5,44^

### A second EGFOA variant adopts a strand-exchanged dimeric architecture

A second EGFOA variant, EGFOA_*Pae*_6186, assembled differently. Instead of populating a stable trimer, this RNA yielded a dimeric state at 3.54 Å resolution in which the two subunits are linked by intermolecular 3′-terminal strand exchange (Figure 4B, G-I and Figure S21-23). This strand displacement generates intermolecular P5 and P7 helices, yielding a compact domain-swapped architecture. Comparison of the monomeric folds from *Vei*_C1A6 and *Pae*_6186 shows that the EGFOA scaffold can be redirected toward at least two distinct quaternary outcomes: a trimer stabilized by distributed tertiary contacts and a dimer stabilized by terminal strand exchange (Figure 4J).

Taken together, the two EGFOA structures show that short RNA assemblies are not restricted to a single favored topology. Even within one family, minor sequence variations, especially with helical rearrangement can redirect the assembly pathway toward fundamentally different non-cage architectures (Figure S24). This behavior parallels a central feature of protein oligomerization,^1^ in which multiple quaternary structures form without requiring major changes in protomer size.^3^

### A 57-nt RNA motif forms an RNA-only filament

We have also asked whether or not relatively short RNA species might assemble in an open-ended manner into filamentous species. The kink-turn (k-turn) is a widespread motif formed in a duplex RNA containing a short unilateral bulge followed by sheared G:A and A:G base pairs.^43^ This forms a kink in the helical axis that can then mediate longer-range contacts, and we have found that k-turns can mediate the formation of defined nano-scale molecular structures.^42^ To this end, we analyzed the 57-nt *Hm* kt7-57. Extended filaments were directly visible in micrographs (Figure 5A). For high-resolution structural analysis, cryo-EM reconstructions of the 6U and 5U variants were obtained at 2.89 Å and 2.72 Å resolution, respectively (Figure S25-27). The resulting maps allowed straightforward atomic model building, with all nucleotides clearly resolved (Figure 5B-C and Figure S28). Notably, the macroscopic dimensions and helical stacking of the *Hm* kt7-57 RNA filament exhibit a marked morphological similarity to naturally occurring biological protein filaments (Figure S29).^7,45,46^

**Figure 5.**
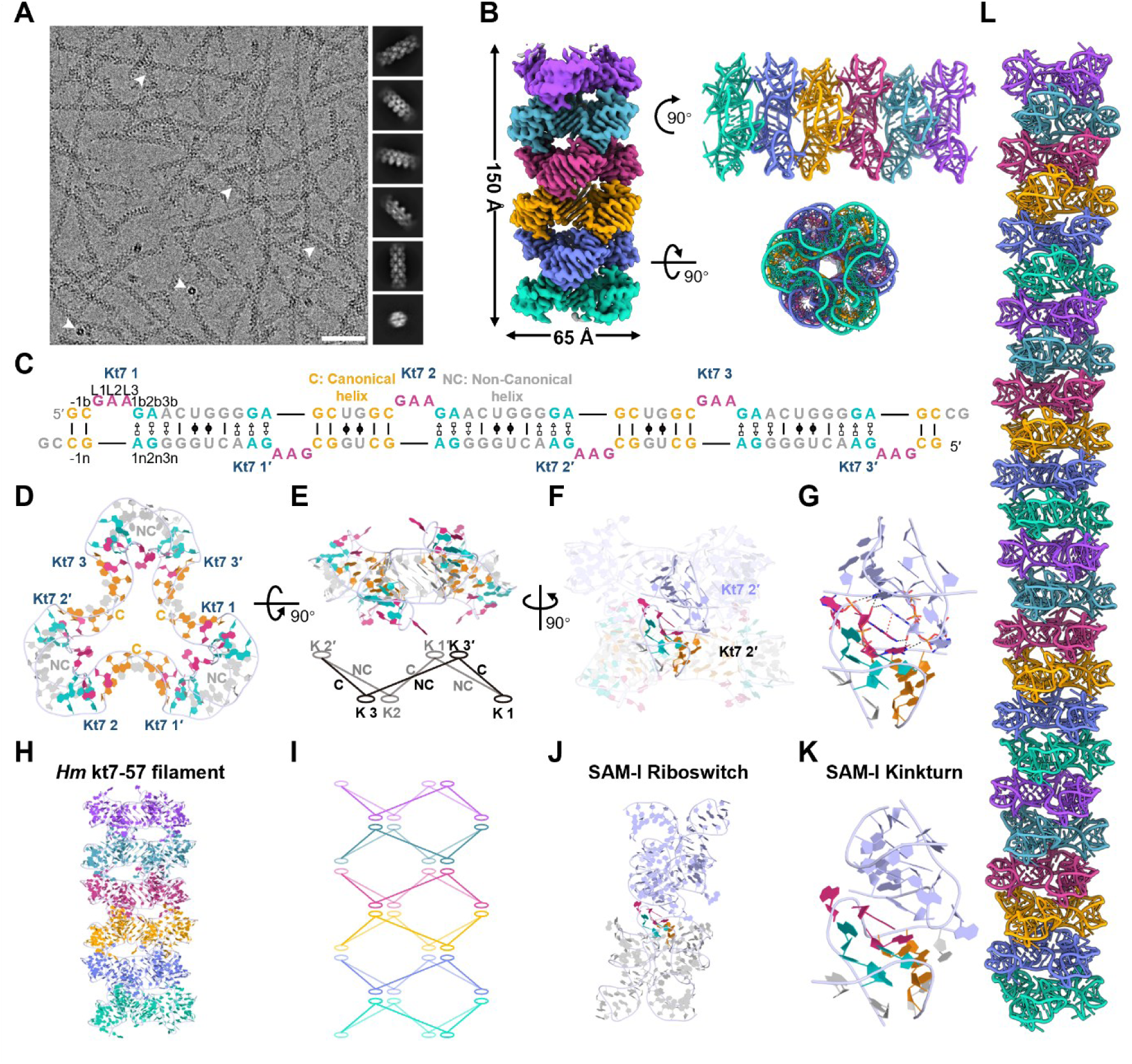
Cryo-EM structure and assembly mechanism of the *Hm* kt7-57 RNA filament. (**A**) Representative cryo-EM micrograph of the *Hm* kt7-57 RNA filaments (white arrowheads). The right panel displays representative 2D class averages detailing the repeating helical features. Scale bar, 40 nm. (**B**) Cryo-EM density map (left) and the corresponding atomic model (right) of the filament, shown in orthogonal views. Notably, the 3D reconstruction of the segmented fibril depicts only a local segment of the RNA fiber, whereas the native fibers exceed 3000 Å in length. (**C**) Sequence and secondary structure diagram of the *Hm* kt7-57-5U RNA. Nucleotide positions are labeled using established kink-turn (k-turn) nomenclature (L1, L2, L3), with the canonical and non-canonical helices denoted as C and NC. (**D**) Top view of the triangular building block formed by the intermolecular assembly of k-turn motifs. (**E**) Side view of the triangular assembly (top) and its topological schematic (bottom). The alternation in direction between the C- and NC-helices generates a quasi-cyclic structure reminiscent of a cyclohexane chair conformation. (**F**) Transparent view of the triangular assembly, highlighting the specific locations of the k-turn interaction modules. (**G**) Detailed atomic view of the k-turn interface. Red dashed lines represent π-π stacking, and black dashed lines indicate hydrogen bonding interactions. (**H**) Overall atomic model of the extended *Hm* kt7-57 RNA filament. (**I**) Schematic representation of the filament assembly, illustrating the continuous, higher-order stacking of the chair-like topological layers. (**J**) Structure of the SAM-I riboswitch. (**K**) Close-up view of the canonical SAM-I k-turn motif. (**L**) Extended atomic model of the *Hm* kt7-57 RNA filament.

This architecture differs fundamentally from the bounded assemblies formed by manA and EGFOA. Rather than forming a closed cage, *Hm* kt7-57 forms a continuous filament extending beyond the field of view in cryo-EM micrographs (Figure 5A). The observation that a 57-nt RNA motif can support such an assembly shows that open-ended propagation, like closed-shell formation, can be generated from a small RNA protomer.

### Reiterative kink-turn interactions drive continuous filament propagation

The repeating unit of the *Hm* kt7-57 filament is a triangular assembly built through intermolecular k-turn interactions (Figure 5D-G). Within each layer, the alternating orientation of the C- and NC-helices generates a quasi-cyclic geometry reminiscent of a cyclohexane chair conformation (Figure 5E). Successive layers then stack through reciprocal docking of upward- and downward-facing k-turn modules, producing a continuous filament by reiterative axial addition of modules by the same interface (Figure 5H,I). This architecture shows how a compact local RNA motif can be amplified into a large open-ended assembly through repeated use of conserved geometric operations and molecular contacts.

The k-turn interface is stabilized by hydrogen bonding and base stacking, with key A-minor interactions at the interface between juxtaposed minor grooves. This is not unique to this filament. Structural comparison with the SAM-I riboswitch and *Hm* kt7-GAAA showed that the same core k-turn geometry is highly conserved across distinct RNA contexts (Figure 5J-K and Figure S30). Thus, the filament does not arise from a de novo engineered interaction principle but from the repetitive arrangement of a compact RNA architectural module that already exists in the natural RNA structure space. A retrospective comparison with the previously determined crystal structure of *Hm* kt7-57-6U further showed that the same basic filament architecture can be identified in the crystal lattice (Figure S31). The new data show that the intermolecular contacts visualized here correspond to a stable assembly solution rather than an artifact of crystal packing.

## DISCUSSION

The main conclusion of this work is that short natural RNAs can access three major architectural regimes of higher-order assembly: a closed shell, defined non-cage oligomers and an open-ended filament (Figure 1A, left). Previous natural RNA-only assemblies had suggested a much narrower picture, dominated by dimers and dihedral multimers built from long RNAs^24–26,28,29^ (Figure 1A, right). Our structures therefore place short-RNA assembly in a broader structural landscape long familiar in protein biology but only rarely recognized in RNA biology (Figure 2D-G, Figure 4K-N and Figure S28).

In structural terms, the importance of the 60-subunit manA assembly lies not simply in its larger stoichiometry, but in the class of organization it represents. Lower-order cyclic or dihedral particles can often be understood as products of local repetition or layered coupling, whereas an icosahedral-like closed shell requires global curvature, coordinated closure and geometric consistency across the entire particle. In this sense, the manA assembly is higher-order not because it is intrinsically ‘better’, but because it enters a more globally constrained regime of organization.^30^ Such geometry also opens a different functional landscape. Relative to lower-order dihedral assemblies, an icosahedral-like shell is more naturally suited to isotropic enclosure, regular surface presentation and compartment-like organization, and therefore offers a more plausible route to cargo sequestration, local concentration and protection of internal components.^47,48^ The significance of the manA structure thus lies in showing that a short natural RNA can encode not only multimerization, but a globally coordinated enclosure-capable architecture.

Intriguingly, while a majority of particles converged into a homogeneous spherical class (∼430 Å in diameter) that were subjected to high-resolution reconstruction, the dataset also revealed larger and morphologically heterogeneous architectures (∼400–750 Å), many of which appeared ellipsoidal or flattened (Figure S32A-C). These features are strikingly reminiscent of the structural polymorphism observed in viral capsids and clathrin-assembled polyhedral coats or pits,^9,48^ suggesting that a subset of these RNAs may organize into higher-order 120-mers or 180-mers.^49^ Notably, we even captured rare instances of assemblies indicative of fission or fusion events (Figure S32D). These observations underscore that short RNAs have the inherent capacity to form complex, potentially cargo-encapsulating particles with architectural properties analogous to those of sophisticated protein-based transport systems.

The EGFOA assemblies reveal a different form of assembly (Figure 4A,B). Rather than forming a shell, they define bounded but open geometries with exposed edges and directional surfaces. In this sense, the boomerang-like trimer is conceptually closer to adaptor-like or hub-like oligomers used for scaffolding and spatial organization than to a compartment.^1^ The strand-exchanged dimer shows that even within the same family, quaternary structure can be reprogrammed through local topological exchange rather than by enlarging the protomer. Together, these states indicate that short RNAs can encode finite anisotropic assemblies whose significance may lie less in enclosure than in generating defined angles, directional connectivity and multivalent interfaces from which new architectures could emerge.

The *Hm* kt7-57 filament extends RNA assembly into a third regime: one-dimensional propagation (Figure 5L). In both shape and assembly pattern, it is closer to a track or framework than to a compartment. Particularly striking is that this behavior is encoded by a 57-nt RNA through repeated reuse of a compact, naturally occurring k-turn interface. Filament formation may therefore represent an especially accessible solution in RNA structure space whenever a local motif imposes directional growth without closure. More broadly, the filament shows that RNA can encode not only bounded objects but also extended linear scaffolds. It is worth noting that we achieved a near-atomic resolution of 2.7 Å for a 57-nt RNA filament, setting a new record for the structural determination of such small, highly repetitive RNA assemblies by cryo-EM.^25,50^

Taken together, the manA cage, the EGFOA oligomers, and the *Hm* kt7-57 filament show that short RNAs can adopt closed, non-caged, and open assemblies in a manner that parallels the major geometric regimes of protein homo-oligomerization (Figure 1). An important implication is that the propensity for complex RNA assembly complexity should not be inferred from transcript length alone. Because the small number of natural RNA-only oligomers characterized so far has largely involved transcripts of several hundred nucleotides, and because small monomeric RNAs have historically been difficult cryo-EM targets, candidate selection has been tacitly biased toward longer RNAs. What matters is not nucleotide count per se, but the presence and geometric arrangement of assembly-competent interaction modules that specify curvature, valency, or axial propagation. More generally, motifs such as kissing loops, GNRA-mediated contacts, and potentially pseudoknot-based interfaces may provide more useful predictors of assembly competence than transcript length alone.^18,20^ In the structures reported here, short RNAs use compact intermolecular motifs to produce cages, defined oligomers, and filaments, indicating that small structured RNAs should be explored not as reduced versions of large assemblies, but as a rich source of distinct quaternary solutions. This shift matters because many structured non-coding RNAs are shorter than 200 nt, and their assembly behavior may therefore be substantially more diverse than currently appreciated.^4140^

The structures also suggest design principles. In protein science, closed shells, finite open scaffolds and linear polymers have repeatedly served as templates for engineered nanostructures. By analogy, the manA 60-mer provides a blueprint for RNA shells with programmable enclosure, EGFOA-like assemblies suggest strategies for encoding directional multivalent scaffolds, and the *Hm* kt7-57 filament shows how compact recurrent motifs can specify persistent one-dimensional frameworks. More broadly, these findings should facilitate the search for additional functional RNA assemblies in nature and inform future efforts in RNA design and structure prediction.^19^

These findings also strengthen the structural plausibility of the RNA world. Classic RNA-world arguments emphasize that RNA could, in principle, unite genotype and phenotype, but an evolving RNA-based system would also have required higher-order organization.^12,13^ Cech highlighted encapsulation as a likely early requirement because compartments can protect nucleic acids, enrich substrates, and couple function to selection.^34^ Our work does not show that the assemblies described here performed such roles in early evolution, nor does it imply that present-day structured RNAs are direct relics of primordial systems. It does show, however, that one such structural capacity—the formation of compartment-like shells by short RNAs—is physically accessible to RNA. The filament and non-cage assemblies similarly suggest that scaffolding and spatial organization could emerge from compact RNA motifs in the absence of proteins. In this way, the demonstration that RNA can form cage and filament-like structures lends further plausibility to the RNA World hypothesis.

## RESOURCE AVAILABILITY

### Lead contact

Further information and requests for resources and reagents should be directed to and will be fulfilled by the lead contact, Lin Huang

### Materials availability

The reagents and materials used in this study are available upon reasonable request.

### Data and code availability

The cryo-EM maps and associated atomic coordinate models have been deposited in the wwPDB OneDep System under EMD accession codes and PDB ID codes of EMD-69109 and 23NP for manA_*pho*_A28C pentamer; EMD-69130 and 23OO for manA_*pho*_A28C decamer; EMD-69111 and 23NR for manA_*pho34*_A767 pentamer; EMD-69112 and 23NS for manA_pho34_A767 decamer; EMD-69755 and PDB 24QD for the focused asymmetric unit of manA_A0F1; EMD-69756 and PDB 24QE for the composite map of manA_A0F1; EMD-69113 and 23NT for manA_67AA monomer; EMD-69108 and 23NO for manA_6FAB monomer; EMD-69115 and 23NU for *Hm* kt7-57-6U; EMD-69116 and 23NV for *Hm* kt7-57-5U; EMD-69107 and 23NN for EGFOA_*Vei*_C1A6 trimer; EMD-69102 and 23ND for EGFOA_*Pae*_6186 dimer.

## Supporting information

Movie S1_manA_A0F1

Movie S3_manA_pho34

## ACKNOWLEDGMENTS

We thank Dr. Keqiong Ye for his valuable discussion. We acknowledge the Cryo-EM System of Instrumental Analysis & Research Center, Sun Yat-sen University, and the Advanced Bio-imaging Technology Platform of Guangzhou National Laboratory for our Cryo-EM data collection. LH is supported by the Natural Science Foundation of China grant 32171191; Guangdong Science and Technology Department (Grant No.2024A1515012594, 2023B1212060013, and 2020B1212030004). ZM is supported by Major Project of Guangzhou Laboratory (Grant No. GZNL2024A01002, GZNL2023A01006, GZNL2025C01007, HWYQ23-003, YW-YFYJ0102), the Natural Science Foundation of China (32270707), and the National Key R&D Programs of China (2025YFE0200600, 2023YFF1204700, 2024YFF1206600). XW is supported by the Guangzhou National Laboratory (Grant No. SRPG22-002) and the Guangdong Pearl River Talent Program (Grant No. 2023QN10Y326).JW is supported by the Natural Science Foundation of Top Talent of SZTU (Grant No. GDRC202418)

## AUTHOR CONTRIBUTIONS

Conceptualization: LH; RNA and Cryo-EM grids preparation: YYR, LZZ, KC; Cryo-EM data collection and processing: YYR, LZZ; Generating RNA atomic coordinates: YYR, LH; Investigation: YYR, LZZ, KC; Visualization: YYR, LZZ, KC, YTX, TTB; Funding acquisition: LH, XPW, ZCM, JW; Project administration: LH; Supervision: JW, ZCM, XPW, LH; Writing – original draft: YYR, LZZ, KC, HL; Writing – review & editing: YYR, LZZ, KC, MXL, YTX, TTB, BWH, BWX, SYH, EW, DMJL, JW, ZCM, XPW, LH. All authors contributed to the preparation of the manuscript.

## DECLARATION OF INTERESTS

The authors declare no competing interests.

## STAR METHODS

### KEY RESOURCES TABLE

| REAGENT or RESOURCE | SOURCE | IDENTIFIER |
| --- | --- | --- |
| **Chemicals, peptides, and recombinant proteins** |  |  |
| ApexHF HS DNA Polymerase | Accurate Biology | AG12202 |
| T7 RNA polymerase | Dr. Lin Huang | N/A |
| NTPs (ATP, UTP, CTP, GTP) | Sangon Biotech | A600020/A620134/A620134/A620570 |
| Spermidine | Sangon Biotech | A600902 |
| **Deposited data** |  |  |
| Cryo-EM map and model of manA_*pho*_A28C pentamer | This paper | EMDB: EMD-69109; PDB: 23NP |
| Cryo-EM map and model of manA_*pho*_A28C decamer | This paper | EMDB: EMD-69130; PDB: 23OO |
| Cryo-EM map and model of manA_*pho34*_A767 pentamer | This paper | EMDB: EMD-69111; PDB: 23NR |
| Cryo-EM map and model of manA_*pho34*_A767 decamer | This paper | EMDB: EMD-69112; PDB: 23NS |
| Cryo-EM map and model of the focused asymmetric unit of manA_A0F1 | This paper | EMDB: EMD-69755; PDB: 24QD |
| Cryo-EM composite map of manA_A0F1 | This paper | EMDB: EMD-69756; PDB: 24QE |
| Cryo-EM map and model of manA_67AA monomer | This paper | EMDB: EMD-69113; PDB: 23NT |
| Cryo-EM map and model of manA_6FAB monomer | This paper | EMDB: EMD-69108; PDB: 23NO |
| Cryo-EM map and model of *Hm* kt7-57-6U filament | This paper | EMDB: EMD-69115; PDB: 23NU |
| Cryo-EM map and model of *Hm* kt7-57-5U filament | This paper | EMDB: EMD-69116; PDB: 23NV |
| Cryo-EM map and model of EGFOA_*Ve*i_C1A6 trimer | This paper | EMDB: EMD-69107; PDB: 23NN |
| Cryo-EM map and model of EGFOA_*Pae*_6186 dimer | This paper | EMDB: EMD-69102; PDB: 23ND |
| **Oligonucleotides** |  |  |
| DNA templates for RNA transcription | This paper | Tables S2 and S3 |
| **Software and algorithms** |  |  |
| cryoSPARC (v4.7.1) | Punjani et al., 2017<sup>51</sup> | https://cryosparc.com/ |
| Topaz | Bepler et al., 2019<sup>52</sup> | https://github.com/tbepler/topaz |
| DeepEMhancer | Sanchez-Garcia et al., 2021<sup>53</sup> | https://github.com/rsanchezgarc/deepEMhancer |
| UCSF ChimeraX | Pettersen et al., 2021<sup>54</sup> | https://www.cgl.ucsf.edu/chimerax/ |
| RNA Composer | Sarzynska et al., 2023<sup>55</sup> | https://rnacomposer.cs.put.poznan.pl/ |
| Coot | Emsley et al., 2010<sup>56</sup> | https://www2.mrc-lmb.cam.ac.uk/personal/pemsley/coot/ |
| PHENIX | Adams et al., 2010<sup>57</sup> | https://phenix-online.org/ |
| MolProbity | Chen et al., 2010<sup>60</sup> | http://molprobity.biochem.duke.edu/ |
| PyMOL | Schrödinger, LLC | https://pymol.org/ |
| **Other** |  |  |
| Superdex 200 Increase 10/300 GL column | Cytiva | 28990944 |
| Centrifugal filters (10 kDa MWCO) | Millipore | 41104916 |
| UltrAuFoil Au 300 mesh, 1.2/1.3 μm grids | Quantifoil | UltrAuFoil R 1.2/1.3 |

### Experimental model and study participant details

#### RNA *In Vitro* Transcription and Purification

To promote efficient transcription initiation, the initial nucleotides of the native sequences were substituted with GGX/ GCG (X = A/U/C/G). DNA templates encoding the T7 promoter upstream of target RNA sequences (Table S2 and 3) were assembled *via* overlap-extension PCR using primer pairs with 15–25 nt terminal complementarity, amplified with ApexHF HS DNA Polymerase (95°C 1 min; 35 cycles: 95°C 30 s, 55°C 30 s, 72°C 30 s; final extension 72°C 5 min), and verified by agarose gel electrophoresis. Large-scale transcription reactions (5 mL) were performed using in-house purified recombinant T7 RNA polymerase (5 mg/mL) in a standardized buffer comprising 40 mM Tris-HCl (pH 7.5), 20 mM MgCl₂, 2 mM spermidine, 4 mM each NTP, 0.25 μM DNA template, 10 mM DTT, and 0.01% Triton X-100, incubated at 37°C for 8–10 h. RNA was precipitated by adding 0.1 volumes of 3 M sodium acetate (pH 5.2) and 1 volume of isopropanol, followed by incubation at –30°C overnight. The RNA pellet was washed with 75% ethanol, air-dried, and dissolved in RNase-free water. Following heat denaturation (95°C, 1 min) in formamide loading buffer, samples were purified on a denaturing polyacrylamide gel containing 7 M urea. The complete RNA transcripts were identified through ultraviolet shadowing, excised, and electroeluted (using the Elutrap Electroelution System, GE Healthcare) in 0.5× Tris-Borate-EDTA (TBE) buffer for 12 h at 200 V at 4°C. Eluted material was ethanol-precipitated, quantified by NanoDrop, and integrity-checked by analytical denaturing urea-PAGE.

#### RNA Refolding and Oligomer Formation

For RNA folding, samples (∼3 mg/mL) were supplemented with 50 mM Tris-HCl (pH 7.5), denatured at 90°C for 3 min, and cooled to room temperature for 10 min. Assembly was initiated by the addition of 10 mM MgCl₂, followed by incubation at 50°C for 30 min. The assembled RNA was then cooled to room temperature for 10 min and stored on ice prior to use.

#### Size-Exclusion Chromatography (SEC)

Refolded RNA samples (500 µL) from six constructs (manA_*pho*_A28C, manA_*pho34*_A767, manA_6FAB, manA_A0F1, EGFOA_*Vei*_C1A6, and *Hm* kt7-57-6U) were subjected to SEC purification on a Superdex 200 Increase 10/300 GL column (Cytiva) using an ÄKTA pure 25 system maintained at 4°C. The column was pre-equilibrated in buffer (20 mM HEPES pH 7.5, 150 mM KCl, 2 mM MgCl₂). Separation was performed at a flow rate of 0.5 mL min⁻¹, and fractions of 0.25 mL were collected throughout the elution profile. Peak fractions corresponding to the main peak were pooled, concentrated using centrifugal filters (10 kDa MWCO), and flash-frozen at –80°C for subsequent cryo-EM grid preparation (Figure S32).

#### Cryo-EM Sample Preparation and Data Acquisition

Sample grids (UltrAuFoil Au 300 mesh, 1.2/1.3 μm) were subjected to glow discharge for 60 s at 15 mA in air. For vitrification, 3 µL of RNA sample was applied to each grid within a Vitrobot Mark IV system (Thermo Fisher Scientific) maintained at 4°C and 100% relative humidity, followed by blotting for 3.5 s prior to rapid plunge-freezing in liquid ethane.

All cryo-EM imaging was performed using 300 kV Titan Krios transmission electron microscopes (Thermo Fisher Scientific) equipped with energy filters and direct electron detectors (K3 Summit camera (Gatan) or Falcon 4i direct electron detector (Thermo Fisher Scientific)). In particular, only the manA_*pho*_A28C dataset was obtained using a 300 kV Titan Krios transmission electron microscope equipped with a spherical aberration corrector. Automated data collection was conducted with EPU software (Thermo Fisher Scientific) using beam-image shift protocols. Movie stacks were recorded at nominal magnification of 130,000×, 165,000×, or 215,000×, corresponding to calibrated pixel sizes of 0.572–0.934 Å. Each exposure comprised 40 frames with a total accumulated dose of 50 e⁻/Å², using defocus values ranging from –0.8 to –1.6 μm. Detailed acquisition parameters for individual datasets are provided in Table S1.

#### Cryo-EM data processing

A total of nine cryo-EM datasets were collected for the manA series, EGFOA series, and *Hm* kt7-57 RNA series. All image processing was performed in cryoSPARC^51^ (v4.7.1). Movie frames were subjected to patch motion correction and patch CTF estimation. Particle picking was initially performed using blob picking, and extracted particles were subjected to 2D classification. Selected particles from good 2D classes were used for ab initio reconstruction and heterogeneous refinement. Templates generated from good 2D class averages or reconstructed 3D maps were subsequently used for template-based particle picking. Particles from blob picking, Topaz picking^52^, and template picking were combined, duplicate particles were removed, and particle subsets were balanced to match the number of good particles identified previously. Seed-facilitated 3D classification ^58^ was performed using heterogeneous refinement models containing good classes as references. Good particle classes were merged and deduplicated, followed by additional rounds of ab initio reconstruction and heterogeneous refinement. High-quality 3D classes were subjected to non-uniform refinement, with or without symmetry constraints depending on the specific dataset, followed by DeepEMhancer^53^ post-processing. For certain datasets, 3D classification, symmetry expansion and particle subtraction were further applied, followed by local refinement and DeepEMhancer optimization. For the OF1 structure, a composite map at 3.34 Å resolution was generated by manually docking monomeric OF1 structures into a low-resolution I-symmetric reference map. Comprehensive cryo-EM data processing workflows and refinement statistics for each structural state are detailed in the supplementary materials (Figure S3, 5, 7, 9, 10, 17, 20, 24, 25 and Table S1).

#### Cryo-EM model building and refinement

Except for the *Hm* kt7-57 RNA series, which used the published crystal structure (PDB: 5G4T)^42^ as the initial model, all other structures were built de novo using guided modeling. Standard A-form RNA duplexes were rigid-body fitted into high-resolution helical density features in ChimeraX, or RNA three-dimensional structures predicted by RNA Composer ^59^ based on secondary structure constraints were used as starting models for density fitting. In Coot^56^, nucleotide types and conformations were manually adjusted to match the corresponding density regions by integrating EM density features, sequence information, and predicted secondary structures. The sequence was progressively extended from both termini, and local conformations were optimized using Real Space Refine Zone until an initial model consistent with the overall density was obtained. This model was subjected to real-space refinement using PHENIX^57^, and the final refined model was validated using MolProbity^60^ for geometric quality assessment. Detailed refinement statistics and validation metrics for each dataset are provided in the Supplementary Table 1. Structure figures were prepared using PyMOL and ChimeraX^54^.

## Notes

### Competing Interest Statement

The authors have declared no competing interest.

