## Supplemental Information for "Structural assemblies for an RNA world"

[1]Guangdong Provincial Key Laboratory of Malignant Tumor Epigenetics and Gene Regulation, Guangdong-Hong Kong Joint Laboratory for RNA Medicine, Medical Research Center, Sun Yat-Sen Memorial Hospital, Sun Yat-Sen University, Guangzhou 510120, China
[2]College of Pharmacy, Shenzhen Technology University, Shenzhen 518118, China
[3]Guangzhou National Laboratory, Guangzhou 510005, China
[4]Wenzhou Institute, University of Chinese Academy of Sciences, Wenzhou 325000, China
[5]Architecture et Réactivité de l'ARN, Université de Strasbourg, Institut de biologie moléculaire et cellulaire du CNRS, Strasbourg, France
[6]Nucleic Acid Structure Research Group, MSI/WTB Complex, The University of Dundee, Dow Street, Dundee DD1 5EH, UK
[7]These authors contributed equally
[8]Lead contact

**Figure S1**

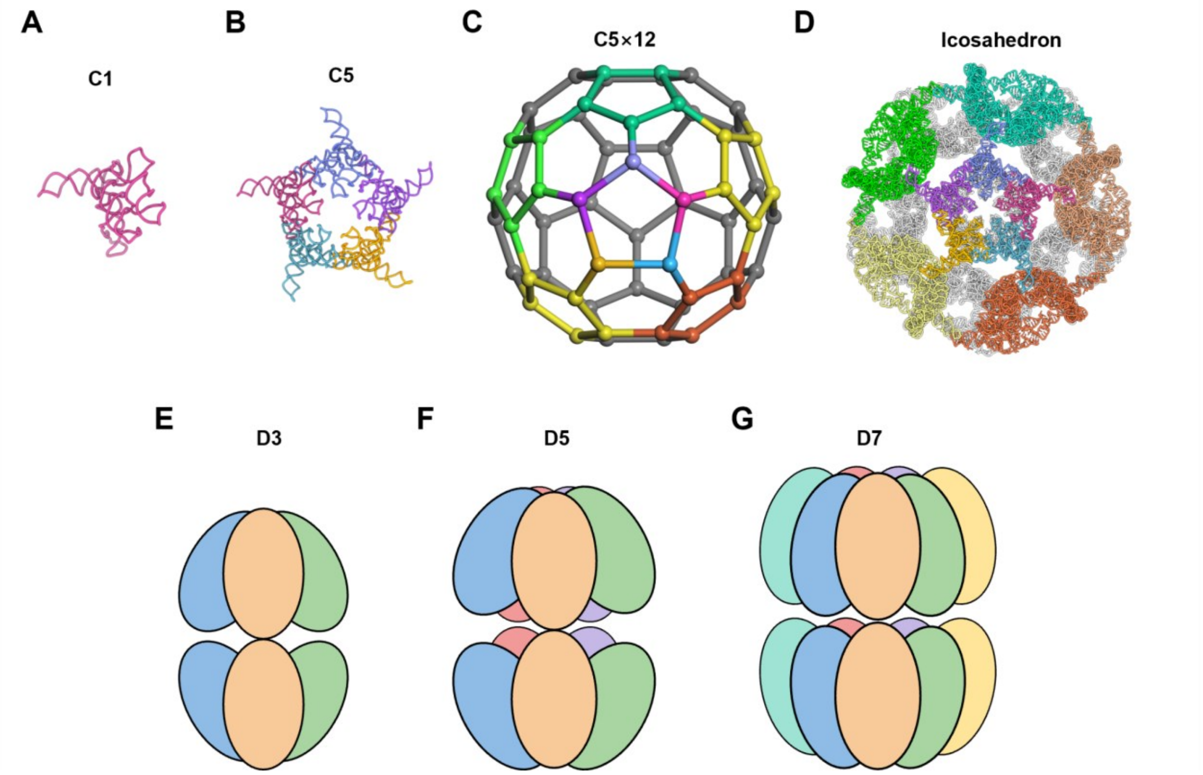


**Figure S1. Architectural distinction between icosahedral and dihedral RNA assembly regimes.** (**A**–**D**) Hierarchical assembly principles of the manA_A0F1 60-mer. The structural progression begins with an asymmetric monomer (C1; **A**), which associates to form a stable pentameric intermediate (C5; **B**). Integration of 12 such pentamers, mapped here onto a fullerene-like geometric framework (**C**), culminates in a fully enclosed spherical cage with strict icosahedral (I) symmetry (**D**). (**E**–**G**) Schematic illustrations of dihedral (D) symmetric architectures, including D3 (**E**), D5 (**F**), and D7 (**G**) geometries. These models represent the prevalent assembly modes of previously reported large natural RNAs (such as ROOL and GOLLD), which predominantly form bipartite, stacked-ring structures. This geometric comparison highlights the more demanding curvature and global closure constraints satisfied by the manA_A0F1 icosahedral shell compared to typical dihedral RNA multimers.

**Figure S2**

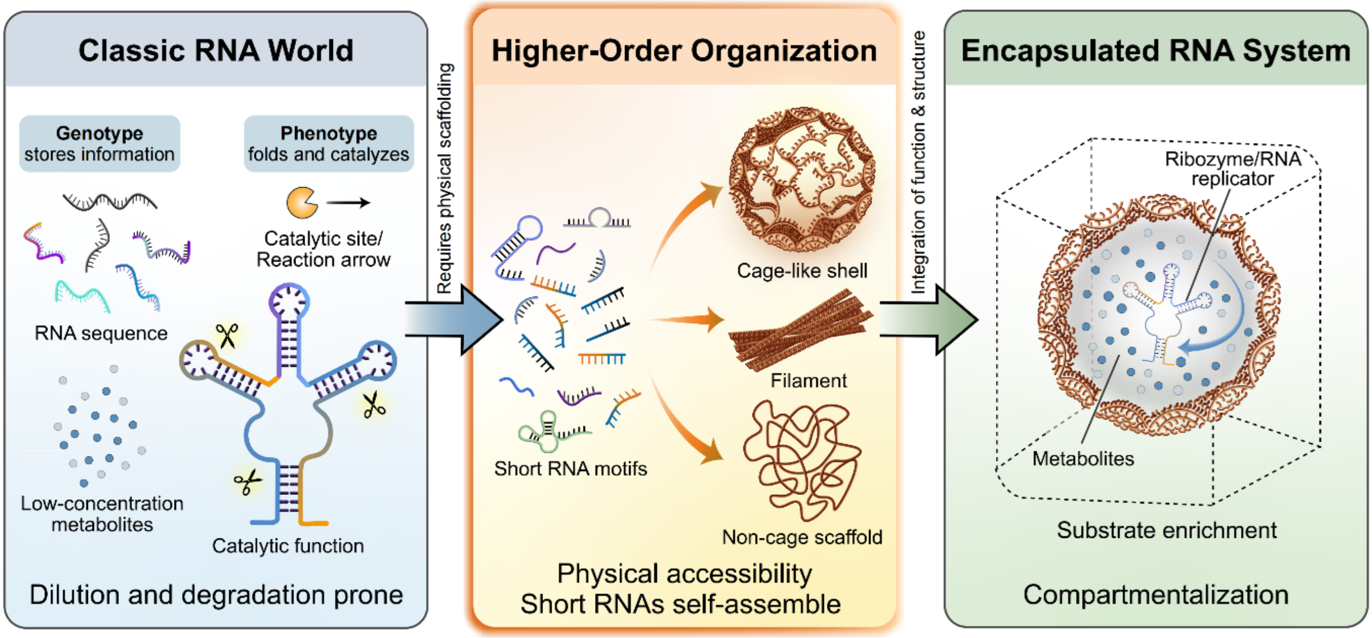


**Figure S2. A structural framework for RNA compartmentalization in the RNA world.** The classic RNA world, shown in the first panel at the left, posits that RNA serves dual roles as both genotype (information storage) and phenotype (catalysis). Higher-order organization is necessary to achieve a programmed and controlled metabolism. As elaborated in this work (second panel), short RNA sequences possess the intrinsic capacity to autonomously self-assemble into diverse macroscopic architectures, including closed cage-like shells, extended filaments, and discrete non-cage scaffolds, besides the large assemblies formed by long RNAs16,17,34. The integration of these structural assemblies with catalytic function represents a critical next frontier (third panel).

**Figure S3**

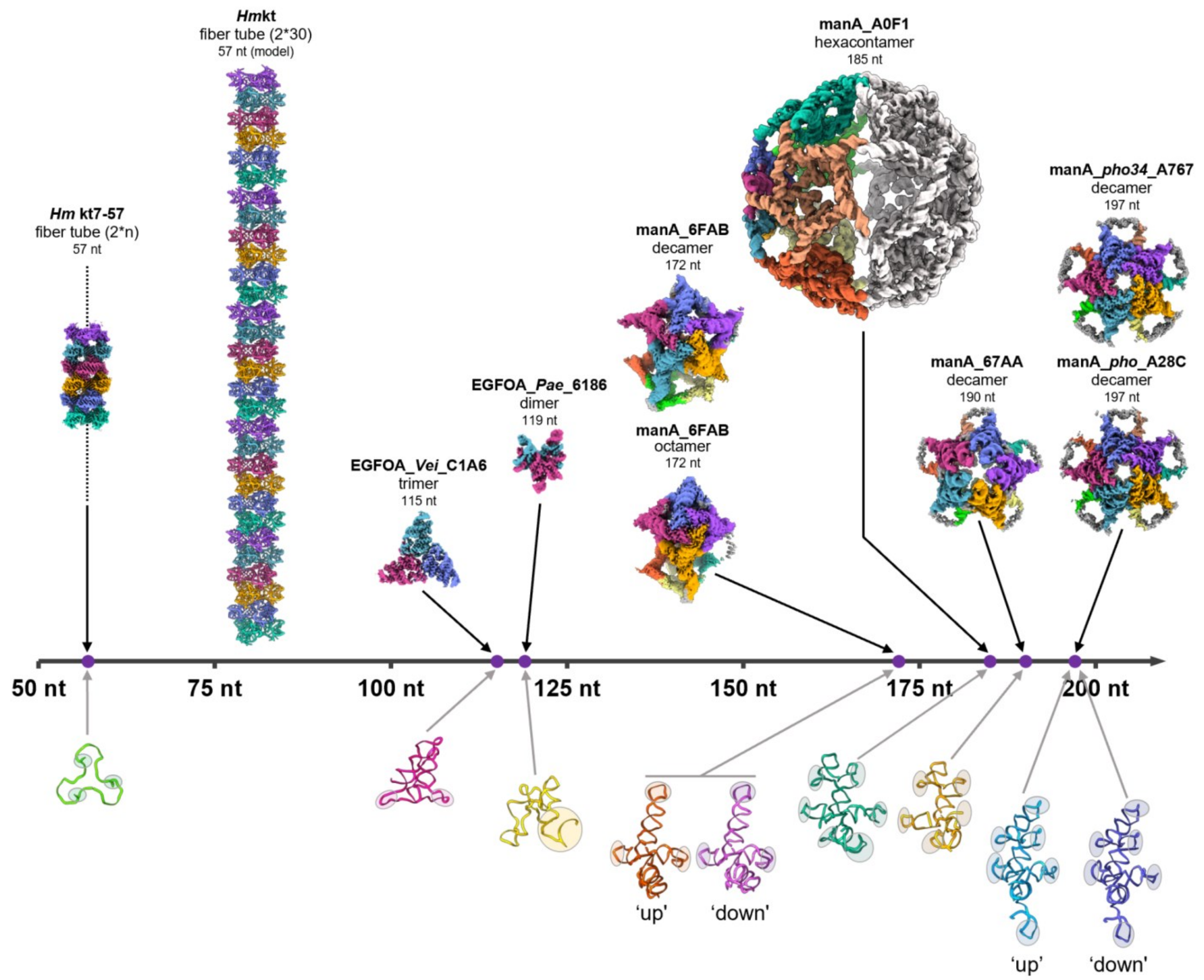


**Figure S3. Structural landscape of the synthesized RNA assemblies and their corresponding monomeric building blocks across nucleotide lengths.** The central axis represents the length of the RNA constructs, ranging from 50 to 200 nucleotides (nt). The higher-order multimeric assemblies presented in the main text (fiber tubes, dimers, trimers, octamers, decamers, and the hexacontamer) are mapped above the axis according to their sequence lengths. Below the axis, the corresponding 3D structures of the monomeric functional units are displayed. Shaded regions explicitly highlight the key structural interfaces that mediate intermolecular interactions and drive self-assembly.

**Figure S4**

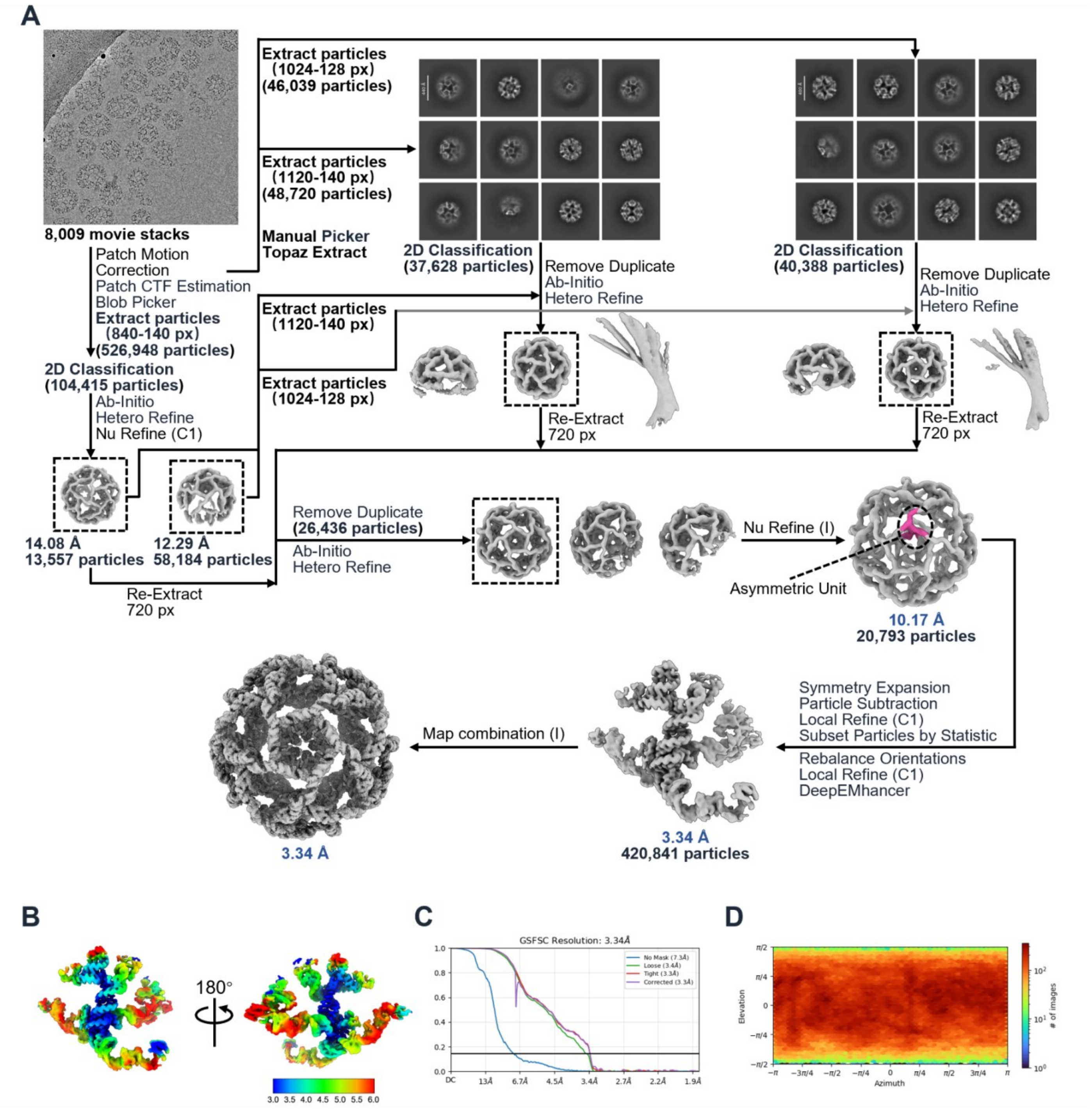


**Figure S4. Cryo-EM data processing workflow for manA_A0F1.** (**A**) Flowchart of the single-particle cryo-EM data processing. (**B**) Local resolution estimation on the cryo-EM map. (**C**) Gold-standard FSC curves for the final reconstruction. (**D**) Euler angle distribution of particles used in the final refinement.

**Figure S5**

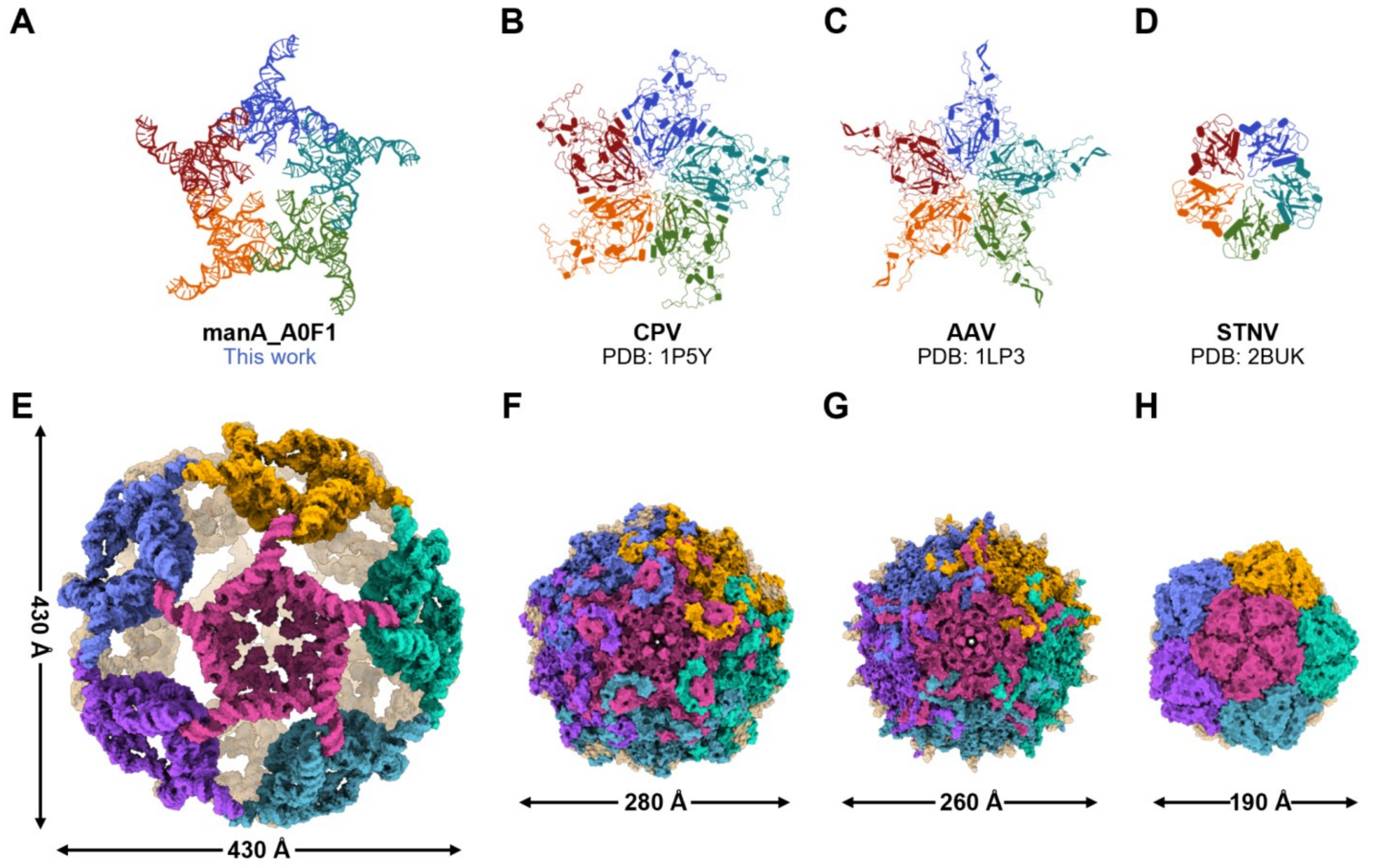


**Figure S5. Structural comparison of the manA_A0F1 RNA 60-mer with naturally occurring T=1 icosahedral viral capsids.** (**A**-**D**) Top views of the fundamental pentameric building blocks for (**A**) the manA_A0F1 RNA assembly (this work), (**B**) Canine Parvovirus (CPV; PDB: 1P5Y)[1], (**C**) Adeno-Associated Virus (AAV; PDB: 1LP3)[2], and (**D**) Satellite Tobacco Necrosis Virus (STNV; PDB: 2BUK)[3]. Structures are shown in cartoon representation with constituent monomers distinctly colored. (**E**-**H**) Surface representations of the corresponding complete 60-mer (T=1) icosahedral assemblies.

**Figure S6**

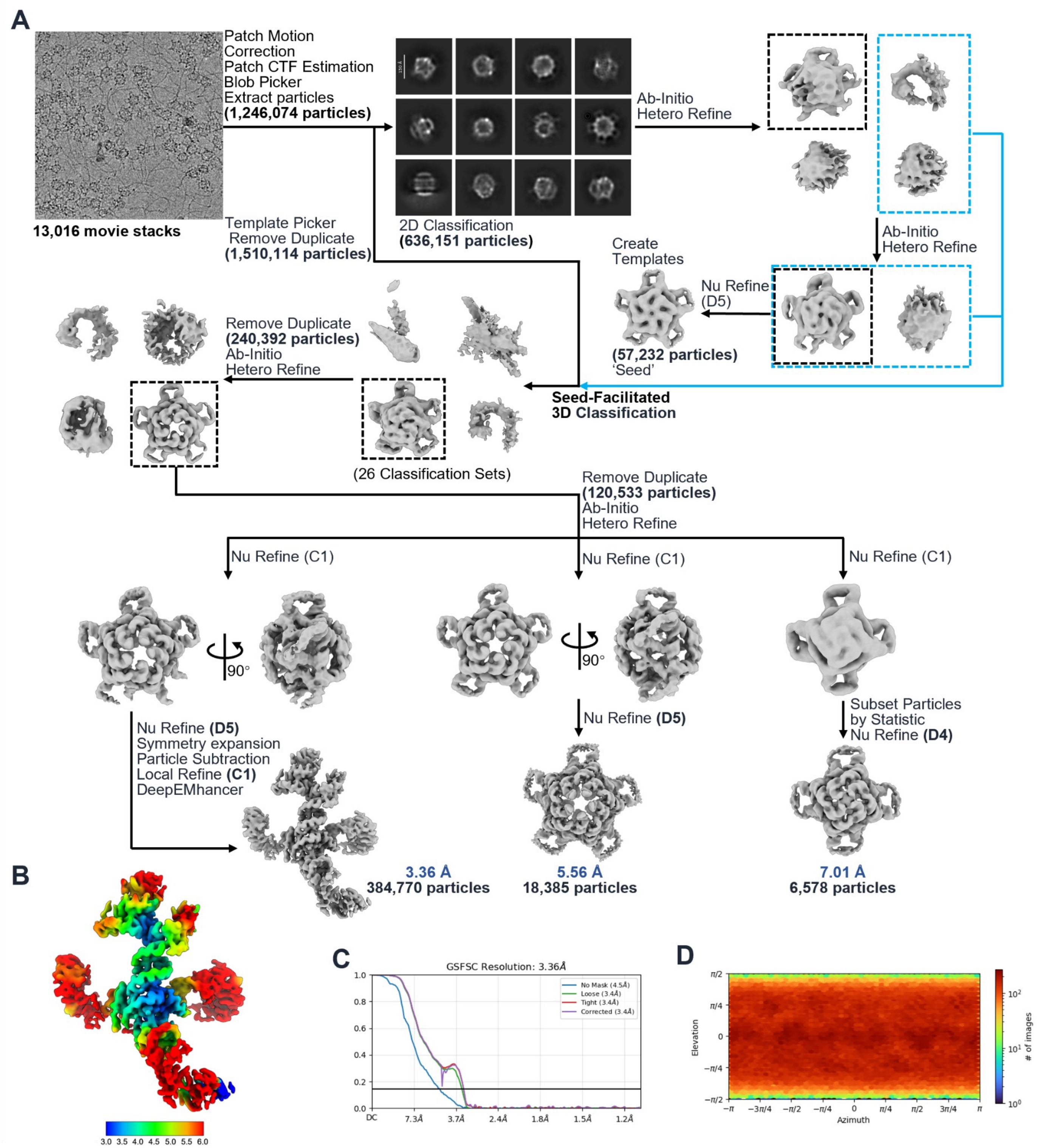


**Figure S6. Cryo-EM data processing workflow for manA_67AA.** (**A**) Flowchart of the single-particle cryo-EM data processing. (**B-F**) Local resolution map (**B**), gold-standard FSC curves (**C**), and Euler angle distribution (**D**).

**Figure S7**

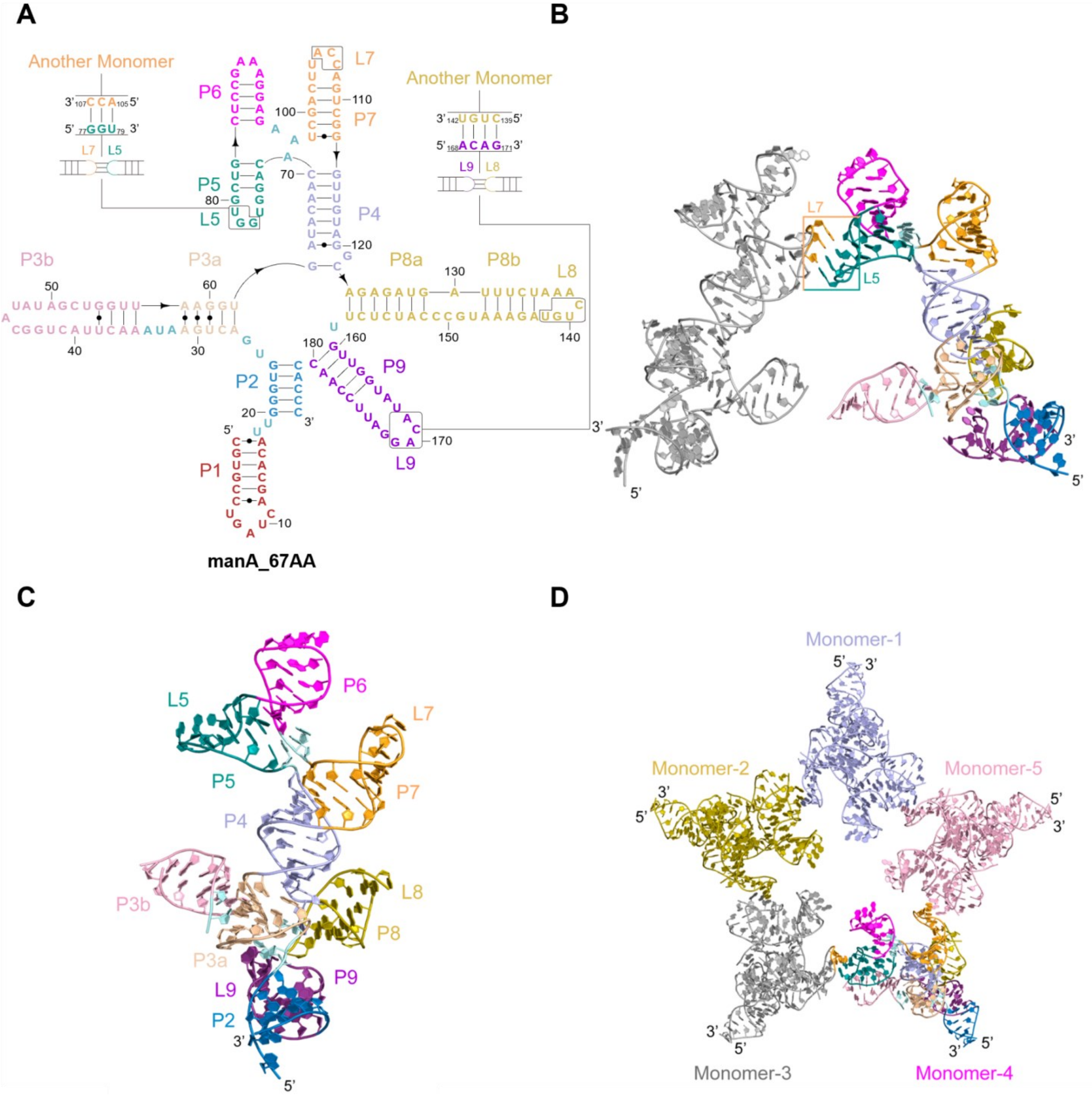


**Figure S7. Structural features and assembly of manA_67AA RNA.** (**A**) Secondary structure of the manA_67AA monomer, with paired regions and loops distinctively colored. Boxed insets show intermolecular kissing-loop interactions. (**B**) Atomic model of the interacting interface. One monomer matches the coloring in (**A**), while the adjacent monomer is gray. The colored box highlights the L5-L7′ interaction. (**C**) 3D model of the manA_67AA monomer, color-coded to match (**A**). (**D**) Overall structure of the manA_67AA pentamer. The five constituent monomers are colored differently to illustrate the assembly.

**Figure S8**

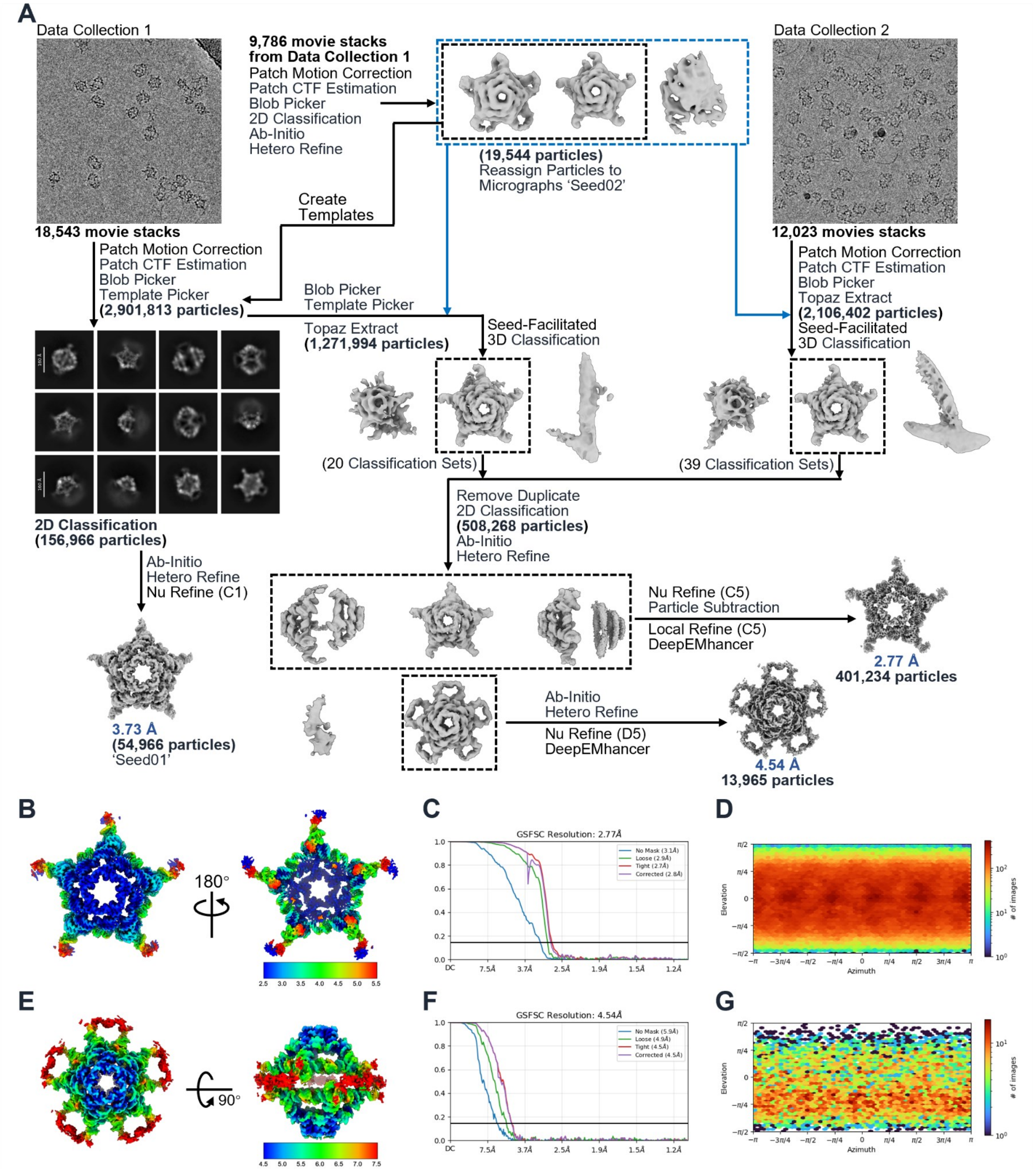


**Figure S8. Cryo-EM data processing workflow for manA_*pho*_A28C.** (**A**) Flowchart of the single-particle cryo-EM data processing. (**B-G**) Local resolution maps (**B**, **E**), gold-standard FSC curves (**C**, **F**), and Euler angle distributions (**D**, **G**).

**Figure S9**

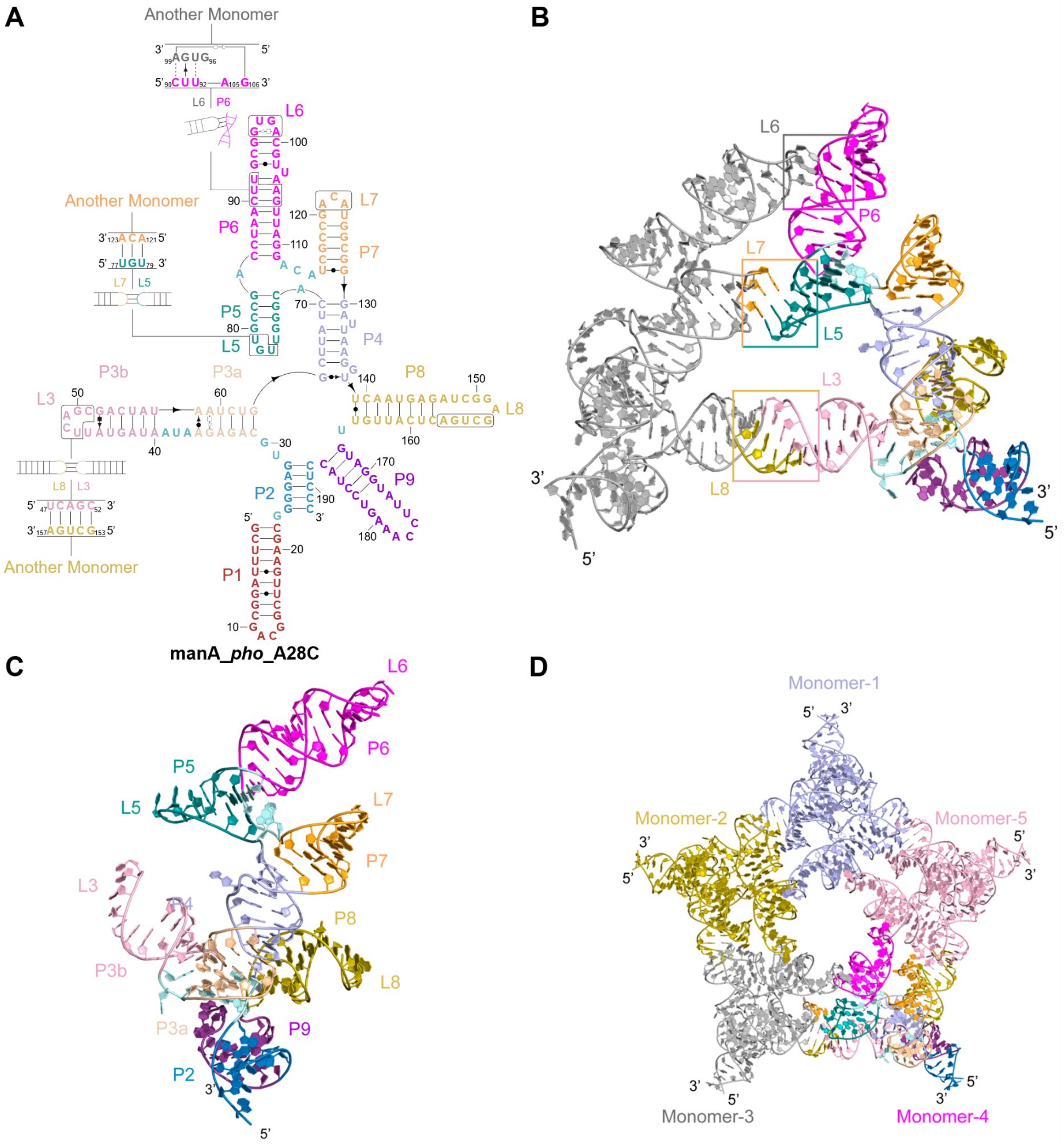

**Figure S9. Structural features and assembly of manA_*pho*_A28C RNA.** (**A**) Secondary structure of the manA_*pho*_A28C monomer, with paired regions and loops distinctively colored. Boxed insets show intermolecular kissing-loop interactions. (**B**) Atomic model of the interacting interface. One monomer matches the coloring in (**A**), while the adjacent monomer is gray. Colored boxes highlight the L6-P6′, L5-L7′, and L3-L8′ interactions. (**C**) 3D model of the manA_*pho*_A28C monomer, color-coded to match (**A**). (**D**) Overall structure of the manA_*pho*_A28C pentamer. The five constituent monomers are colored differently to illustrate the assembly.

**Figure S10**

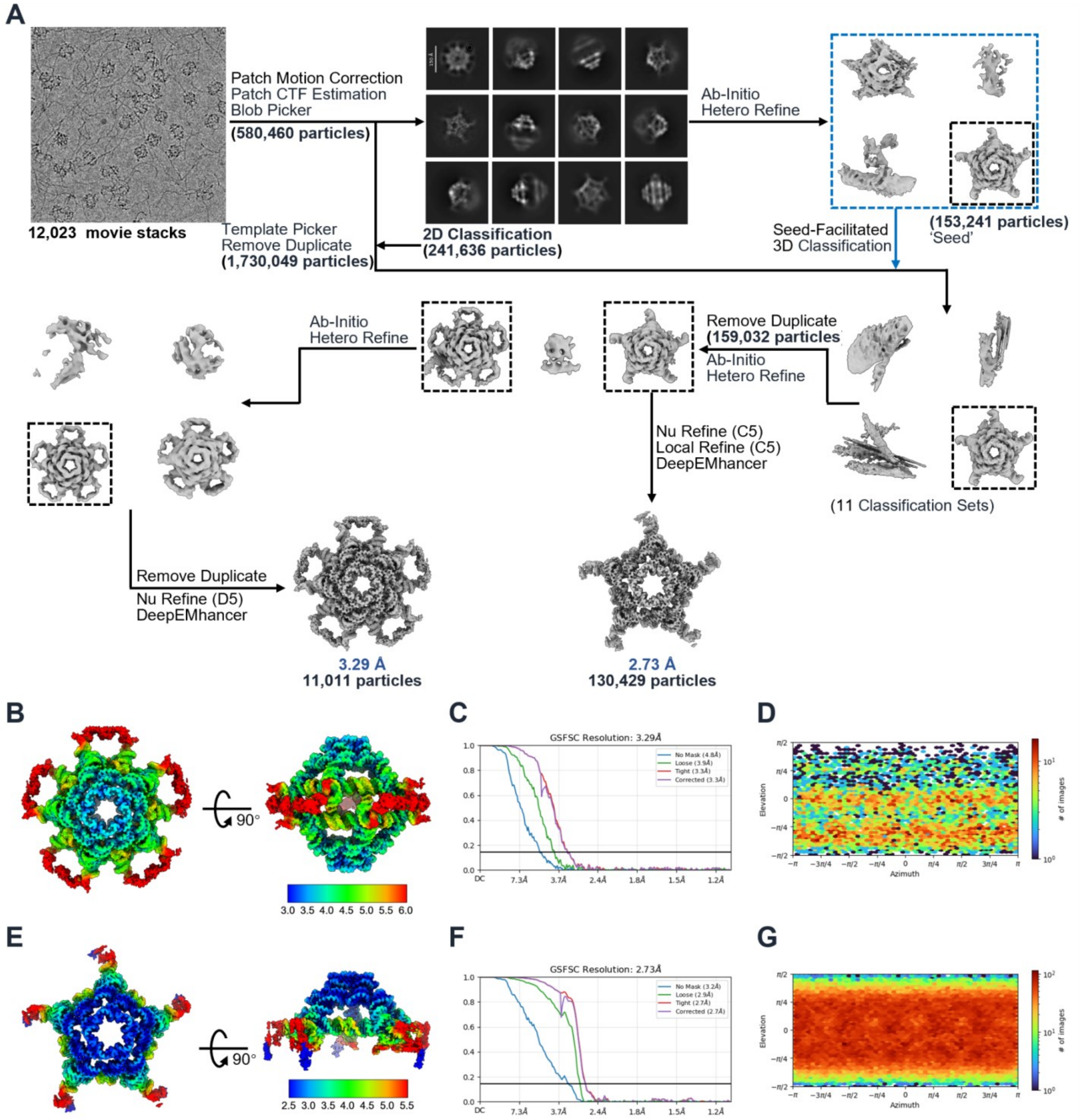


**Figure S10. Cryo-EM data processing workflow for manA_*pho34*_A767.** (**A**) Flowchart of the single-particle cryo-EM data processing. (**B-G**) Local resolution maps (**B**, **E**), gold-standard FSC curves (**C**, **F**), and Euler angle distributions (**D**, **G**).

**Figure S11**

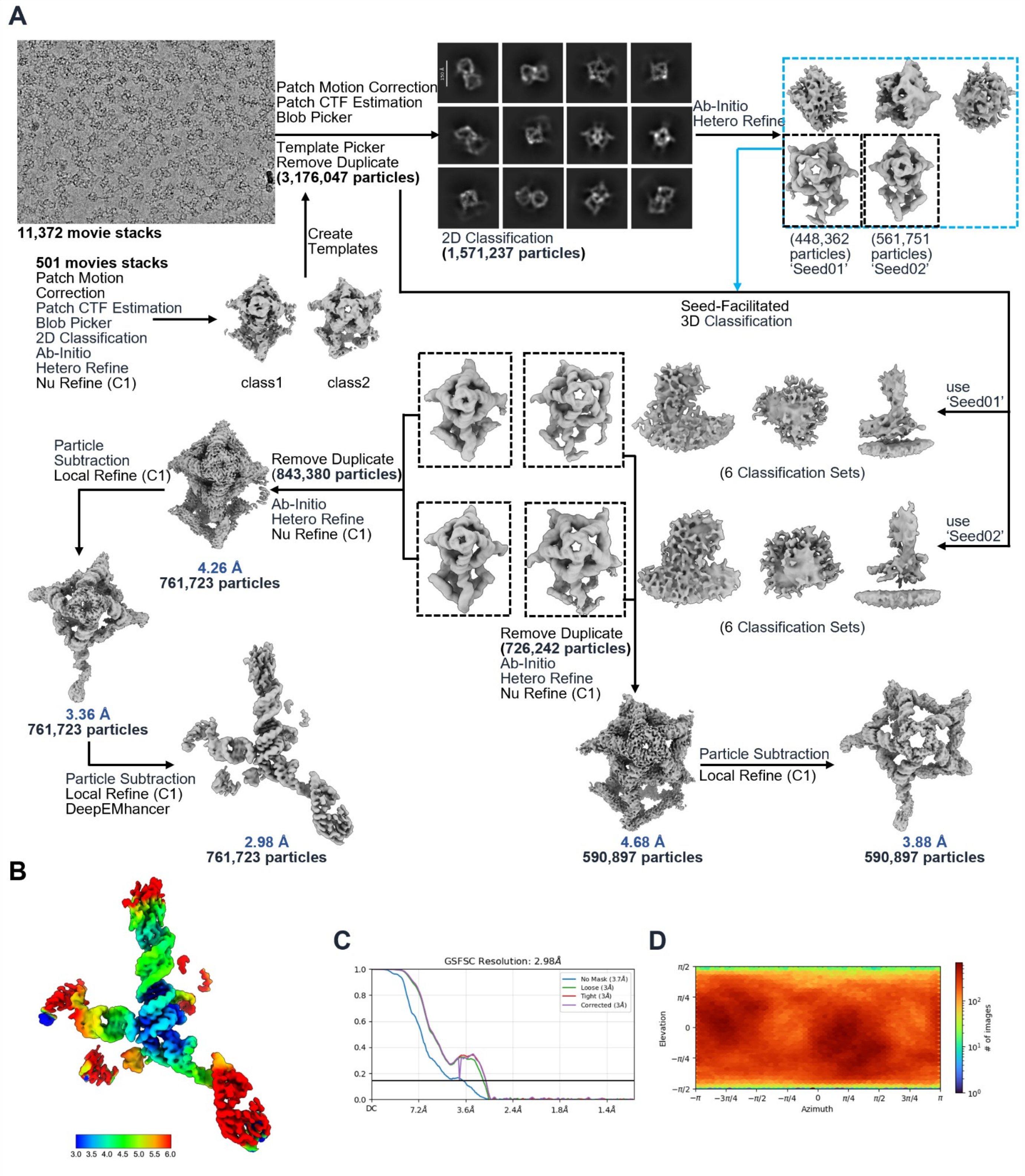


**Figure S11. Cryo-EM data processing workflow for manA_6FAB.** (**A**) Flowchart of the single-particle cryo-EM data processing. (**B**-**D**) Local resolution map (**B**), gold-standard FSC curves (**C**), and Euler angle distribution (**D**).

Figure S12

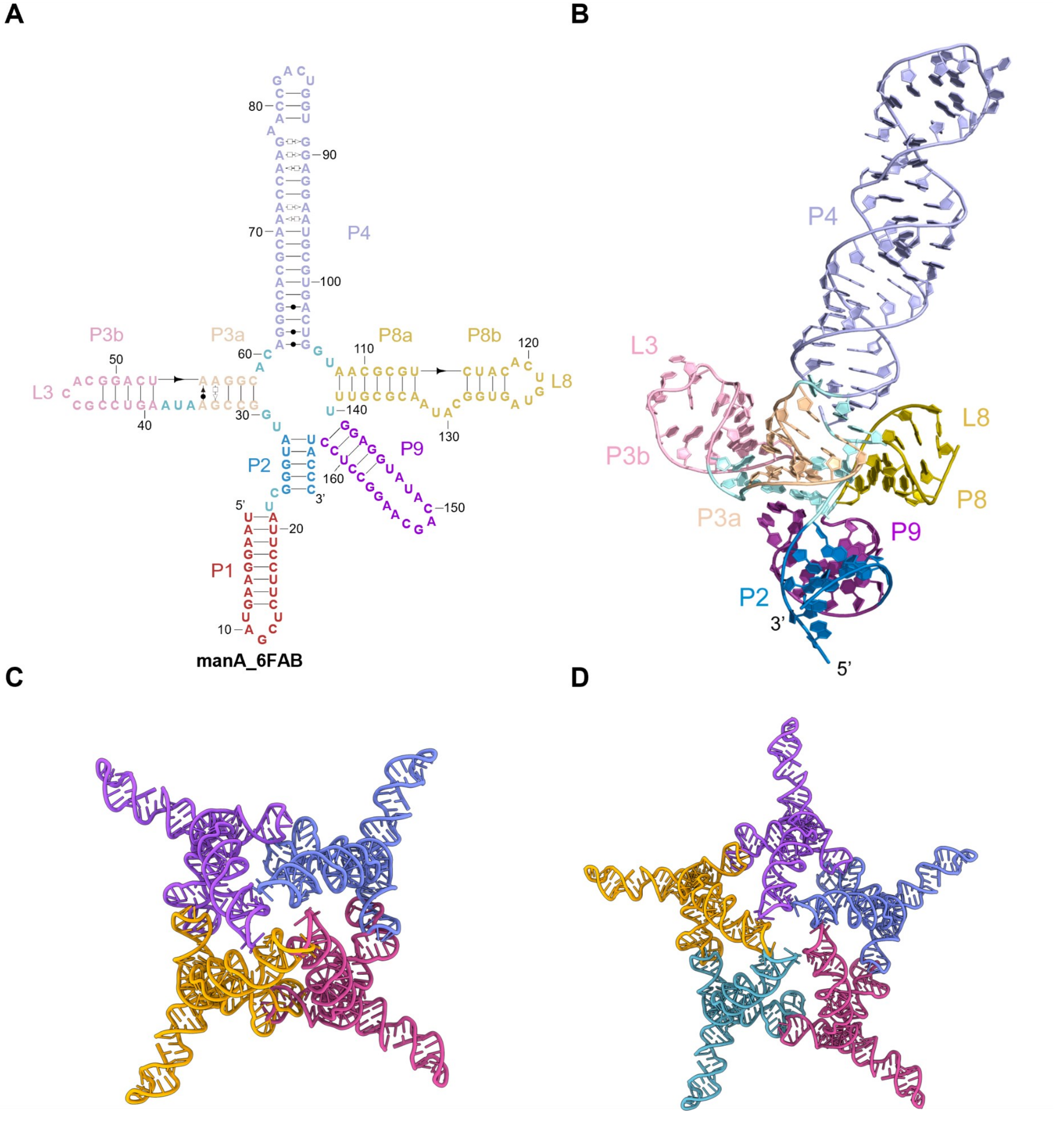


**Figure S12. Structural features and assembly polymorphism of manA_6FAB RNA.** (**A**) Secondary structure of the manA_6FAB monomer, with paired regions and loops distinctively colored. (**B**) 3D model of the manA_6FAB monomer, color-coded to match the secondary structure diagram in (**A**). (**C**) Overall structure of the manA_6FAB tetramer. The four constituent monomers are colored differently to illustrate the assembly. (**D**) Overall structure of the manA_6FAB pentamer. The five constituent monomers are colored differently to illustrate the higher-order assembly.

### Figure S13

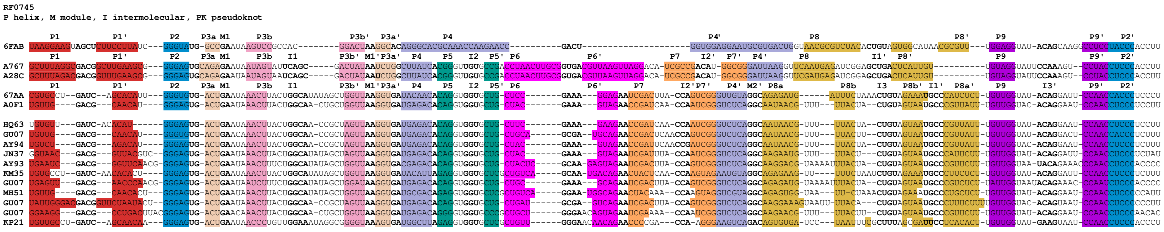

**Figure S13. Structure-based sequence alignment of the manA RNA family.** Multiple sequence alignment comprising the experimentally characterized manA variants (6FAB, A767, A28C, 67AA, and A0F1) and representative homologous sequences from the RF0745 family. Conserved secondary and tertiary structural elements are highlighted with color-coded boxes, corresponding to the structural models presented in previous figures. While A767, A28C, 67AA, and A0F1, as well as the majority of the sequences in RF0745 contain a four-way junction (P4-P5-P6-P7), 6FAB does not. Because P6 is longer in A767 and A28C than in 67AA and A0F1, the quaternary contacts formed are not identical. In A767 and A28C, there are three intermolecular contacts, named I1 (loops L5-L7), I2 (loops L3-L8), and I3 (GNRA-helical Watson-Crick pairs in the middle of P6). Instead, in 67AA and A0F1, there are only two, one is common I1, while I2 shares a common loop (L8) but targets L9 instead of L3). However, the intramolecular modules formed by non-Watson-Crick pairs within helices, M1 (between P3a and P3b) and M2 (at the base of P4, the entry of the four-way junction), are maintained.

**Figure S14**

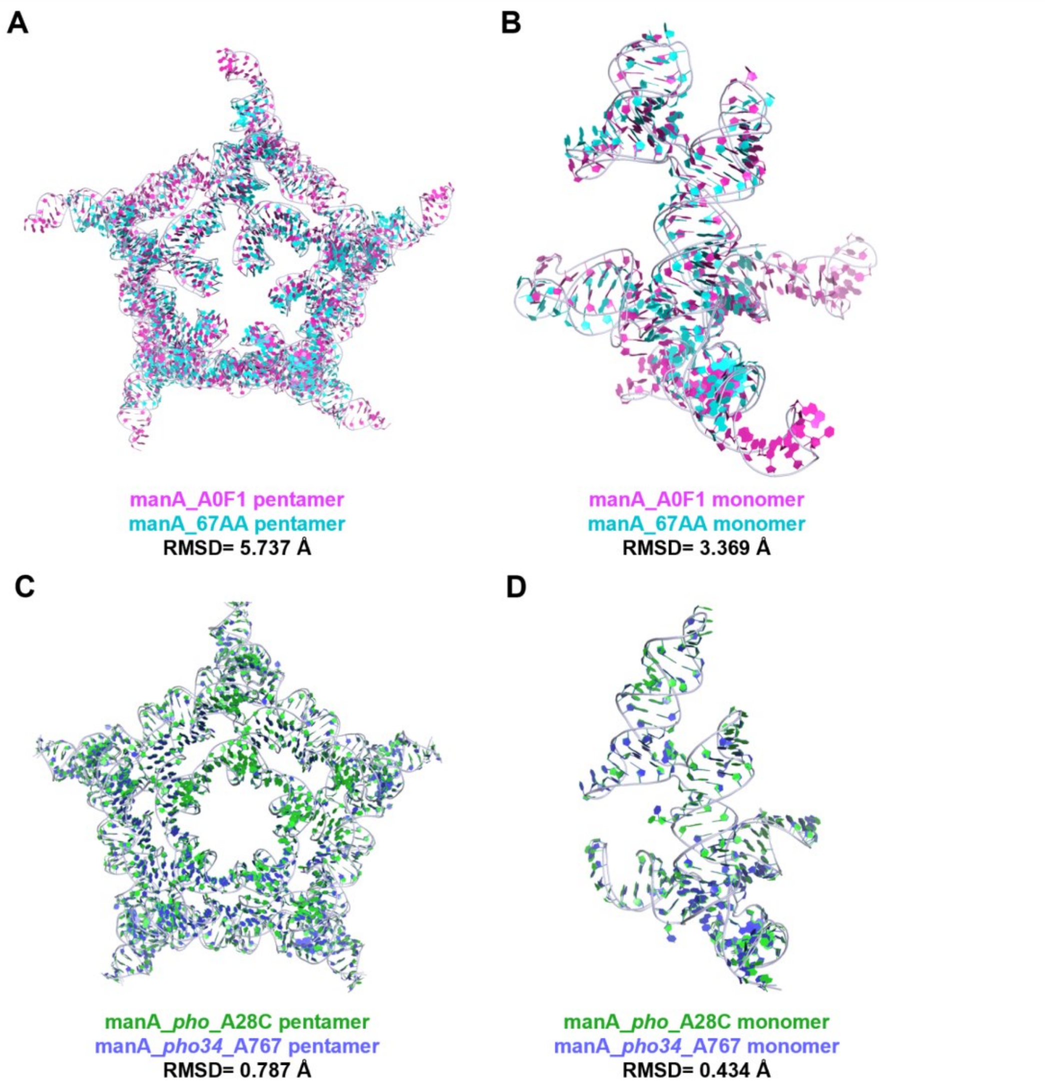


**Figure S14. Structural superpositions of the manA RNA variants.** (**A**) Structural alignment of the manA_A0F1 (magenta) and manA_67AA (cyan) pentamers, showing an overall root-mean-square deviation (RMSD) of 5.737 Å. (**B**) Superposition of the corresponding manA_A0F1 and manA_67AA monomers, with an RMSD of 3.369 Å. (**C**) Structural alignment of the manA_pho_A28C (green) and manA_pho34_A767 (purple) pentamers, yielding a highly conserved RMSD of 0.787 Å. (**D**) Superposition of the corresponding manA_*pho*_A28C and manA_*pho34*_A767 monomers, with an RMSD of 0.434 Å.

#### Figure S15

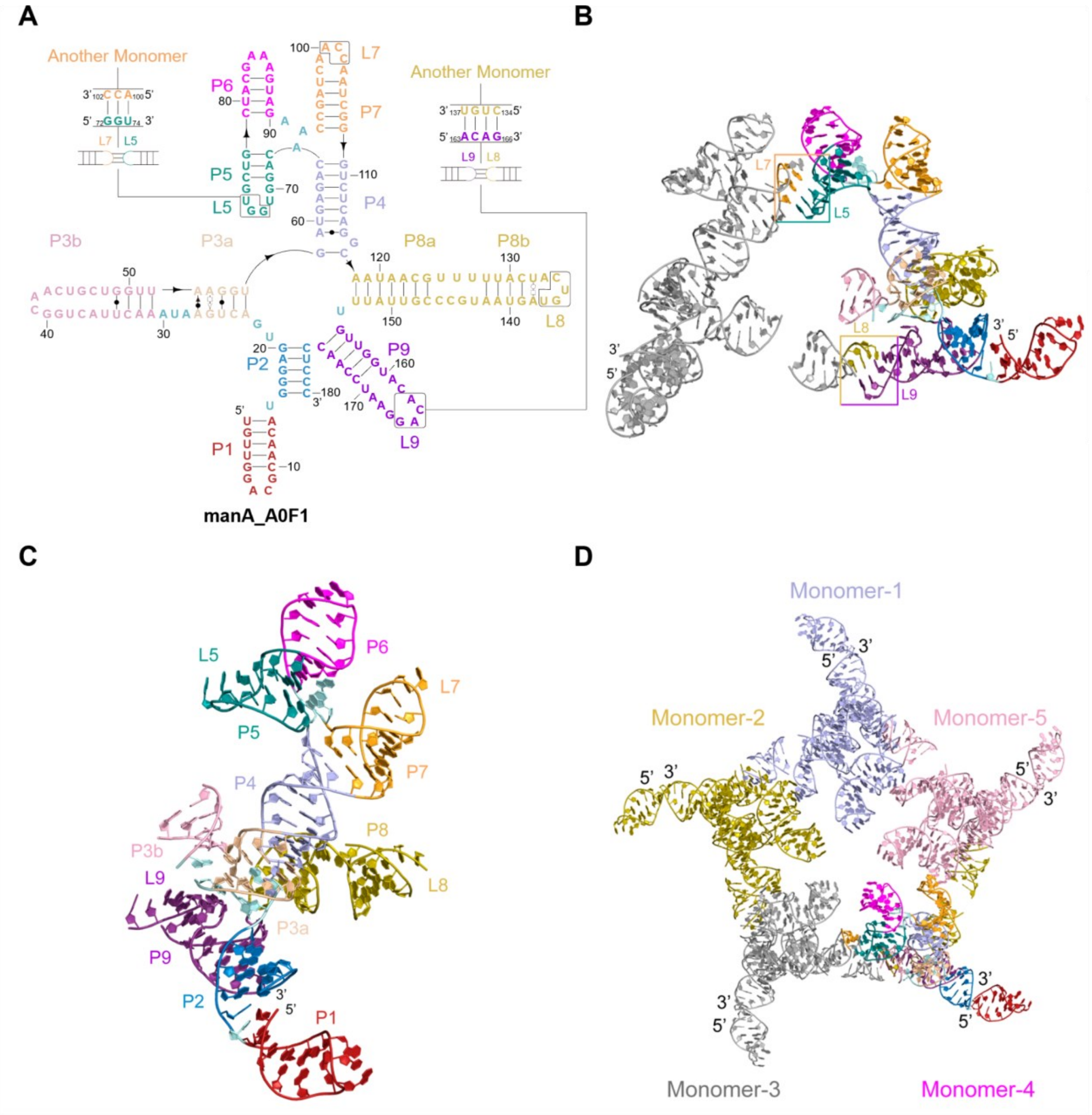


**Figure S15. Structural features and assembly of manA_A0F1 RNA.** (**A**) Secondary structure of the manA_A0F1 monomer, with paired regions and loops distinctively colored. Boxed insets show intermolecular kissing-loop interactions. (**B**) Atomic model of the interacting interface. One monomer matches the coloring in (**A**), while the adjacent monomer is gray. Colored boxes highlight the L5-L7′ and L8-L9′ interactions. (**C**) 3D model of the manA_A0F1 monomer, color-coded to match (**A**). (**D**) Overall structure of the manA_A0F1 pentamer. The five constituent monomers are colored differently to illustrate the assembly.

Figure S16

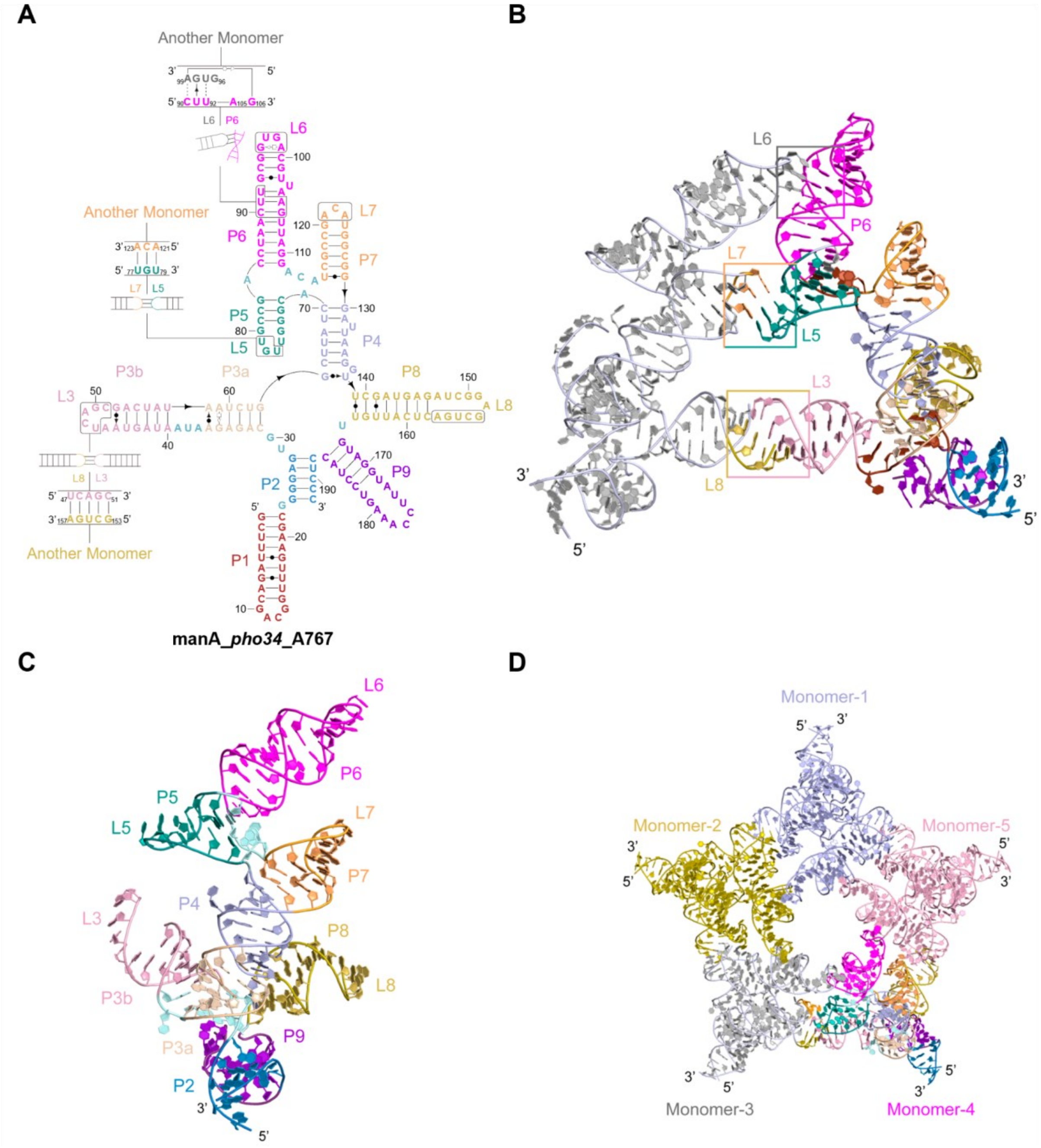


**Figure S16. Structural features and assembly of manA_*pho34*_A767 RNA.** (**A**) Secondary structure of the manA_*pho34*_A767 monomer, with paired regions and loops distinctively colored. Boxed insets show intermolecular kissing-loop interactions. (**B**) Atomic model of the interacting interface. One monomer matches the coloring in (**A**) (**C**) 3D model of the manA_*pho34*_A767 monomer, color-coded to match (**A**). (**D**) Overall structure of the manA_*pho34*_A767 pentamer. The five constituent monomers are colored differently to illustrate the assembly.

**Figure S17**

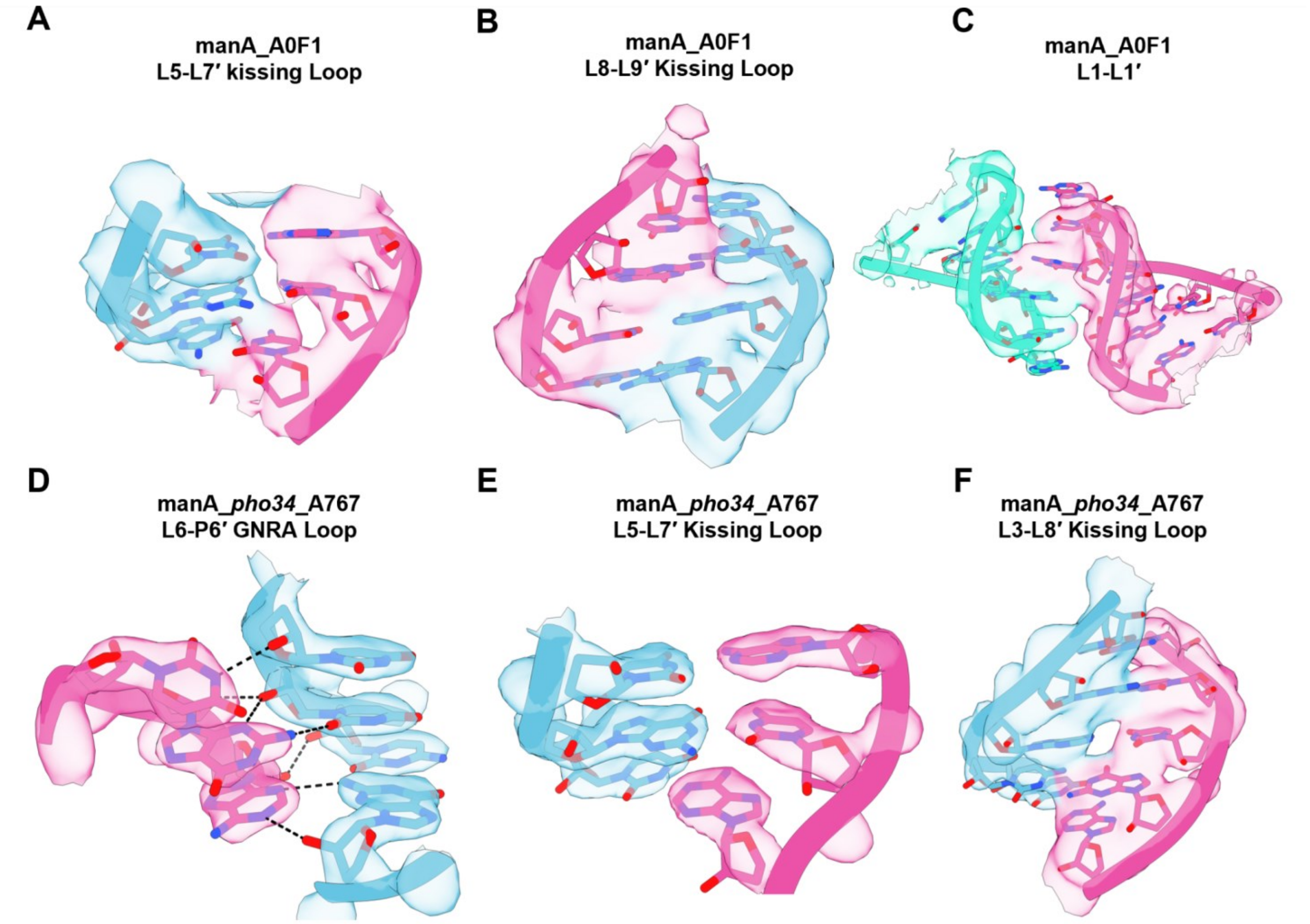


**Figure S17. Cryo-EM density maps validating the key intermolecular interactions.** Detailed views of the structural interfaces supplement the atomic models presented in the main text. The corresponding cryo-EM densities are shown as transparent surfaces to validate the atomic modeling. RNA backbones and bases are shown in stick representation, with the two interacting monomers colored in cyan and magenta, respectively. (**A**-**C**) Close-up views of the L5-L7′ (**A**), L8-L9′ (**B**), and L1-L1′ (**C**) interactions mediating the assembly of manA_A0F1. (**D** to **F**) Close-up views of the intermolecular interfaces in manA_*pho34*_A767, detailing the L6-P6′ GNRA loop interaction with dashed lines indicating putative hydrogen bonds (**D**), the L5-L7′ kissing loop (**E**), and the L3-L8′ kissing loop (**F**).

Figure S18

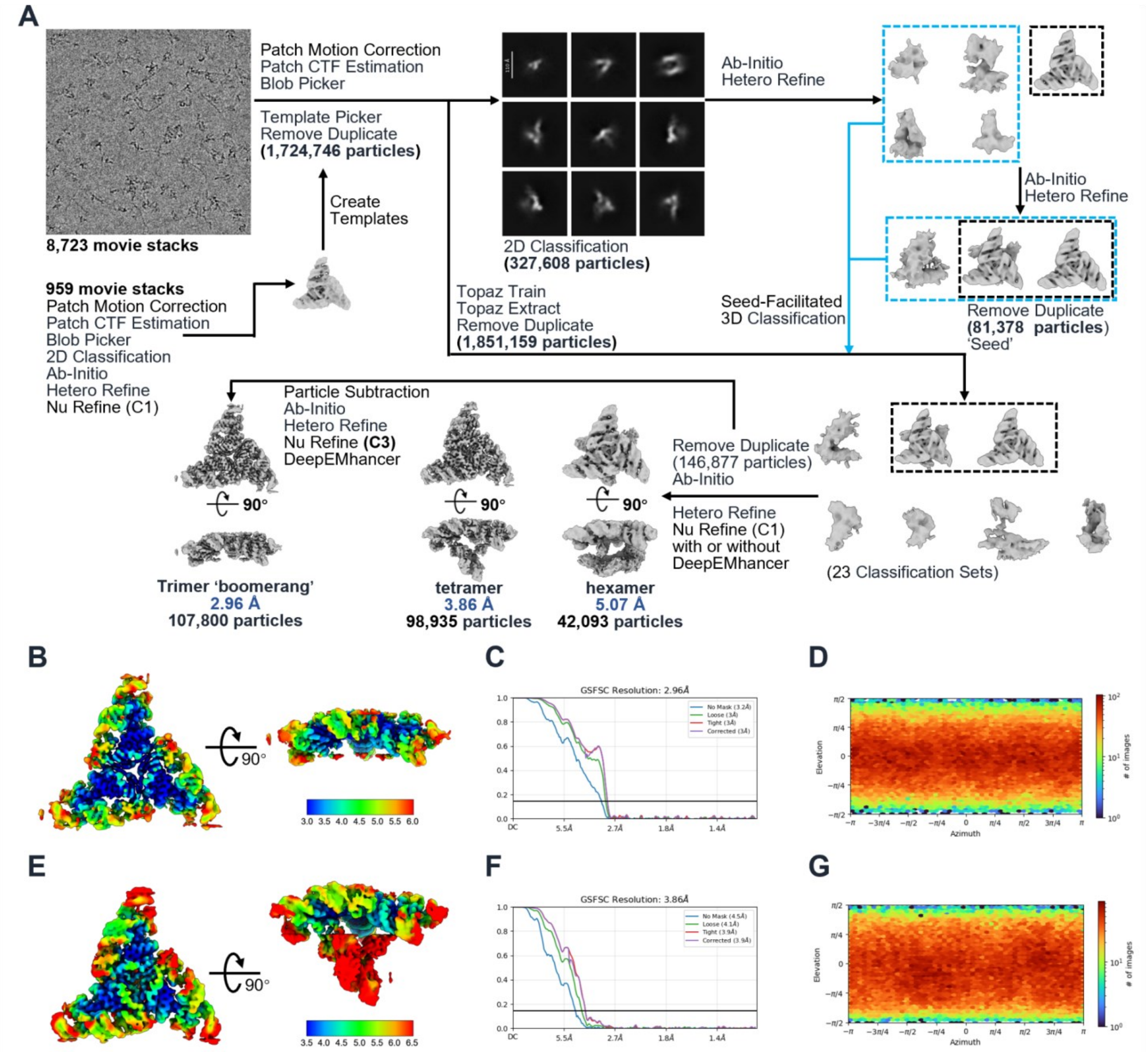


**Figure S18. Cryo-EM data processing workflow for EGFOA_*Vei*_C1A6.** (**A**) Flowchart of the single-particle cryo-EM data processing. (**B**-**G**) Local resolution maps (**B**, **E**), gold-standard FSC curves (**C**, **F**), and Euler angle distributions (**D**, **G**).

**Figure S19**

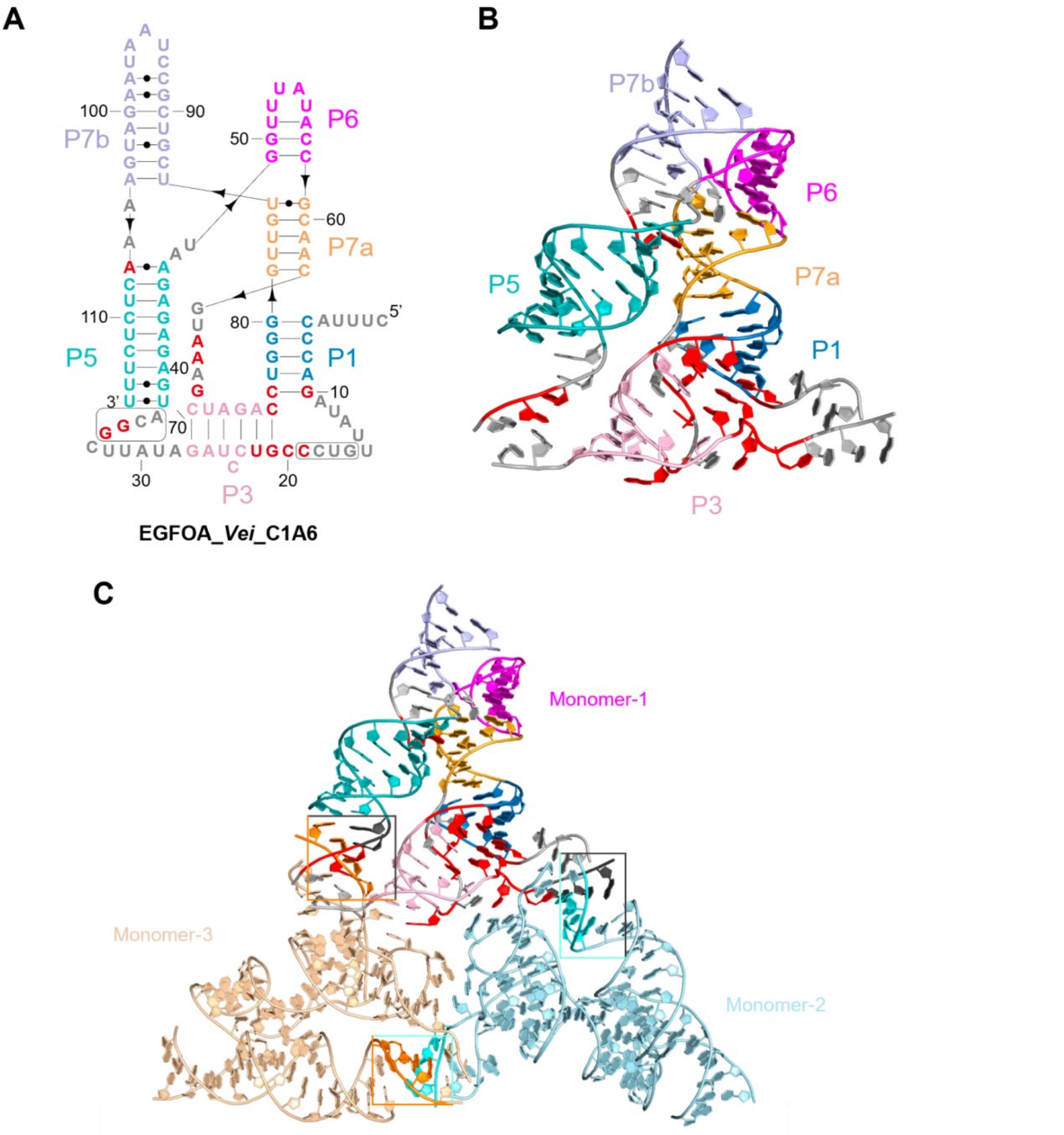


**Figure S19. Structural features and assembly of EGFOA_Vei_C1A6 RNA.** (**A**) Secondary structure of the EGFOA_*Vei*_C1A6 monomer, with paired regions distinctively colored. Highly conserved nucleotides are explicitly highlighted in red. (**B**) 3D atomic model of the EGFOA_*Vei*_C1A6 monomer. Structural elements are color-coded to match the secondary structure diagram in (**A**), with the conserved bases identically mapped in red. (**C**) Overall architecture of the EGFOA_*Vei*_C1A6 trimer. The three constituent monomers (Monomer-1 to Monomer-3) are colored differently to illustrate the 'boomerang' assembly, with dashed boxes highlighting the key intermolecular interfaces.

**Figure S20**

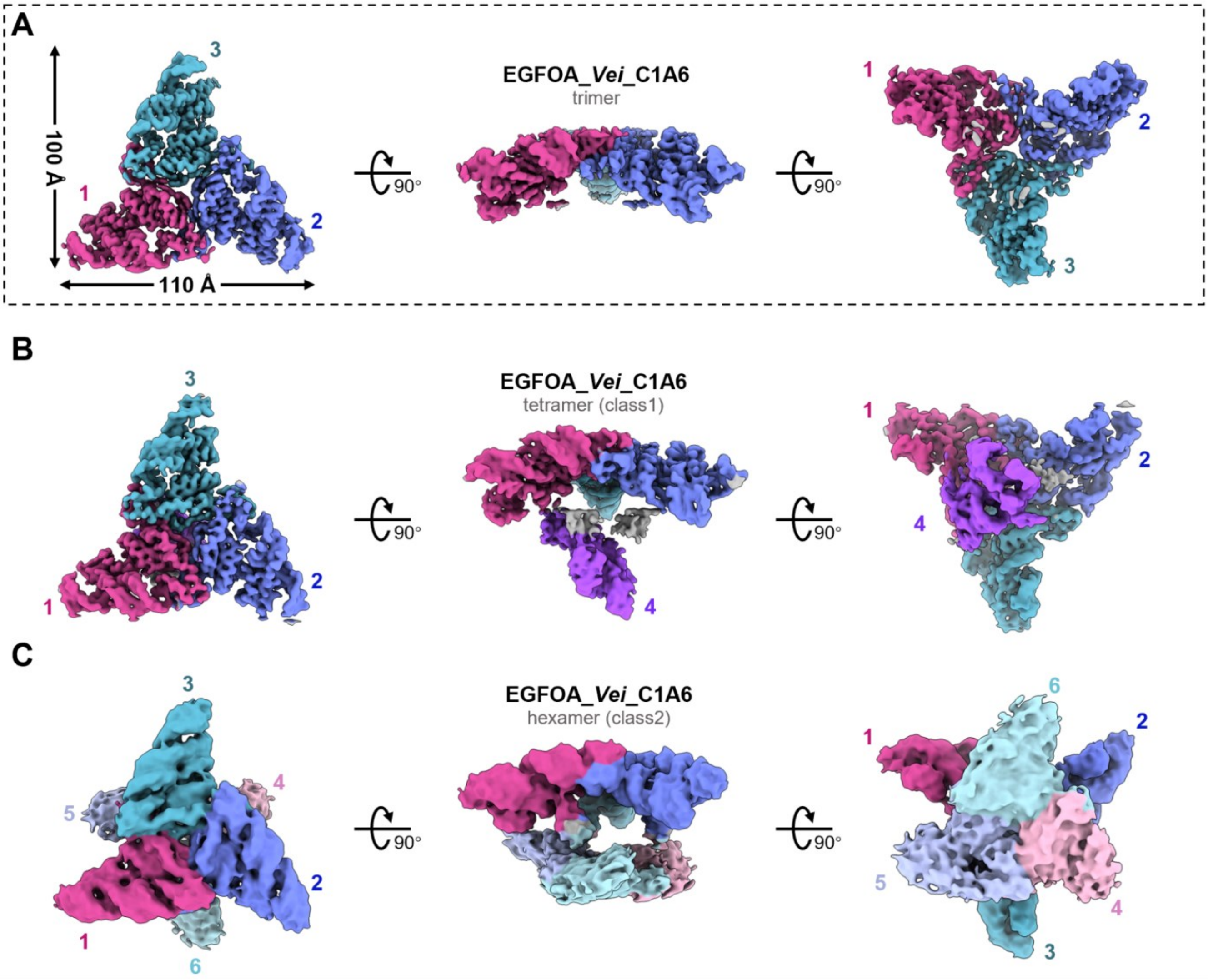


**Figure S20. Structural polymorphism and dynamic assembly states of the EGFOA_*Vei*_C1A6 RNA.** Cryo-EM density maps illustrating the diverse oligomeric states captured during the dynamic assembly process. Each state is shown in three orthogonal views (rotated by 90°), with individual monomers distinctly colored and numbered to trace the structural progression. (**A**) The fundamental trimeric assembly, with its overall dimensions (~100 Å × 110 Å) indicated. (**B**) The tetrameric assembly state (class 1), highlighting the recruitment of an additional monomer (labeled 4, purple) to the base trimeric core. (**C**) The higher-order hexameric assembly state (class 2) demonstrates the expansion of the complex through the incorporation of three additional monomers (labeled 4, 5, and 6) onto the trimeric core.

**Figure S21**

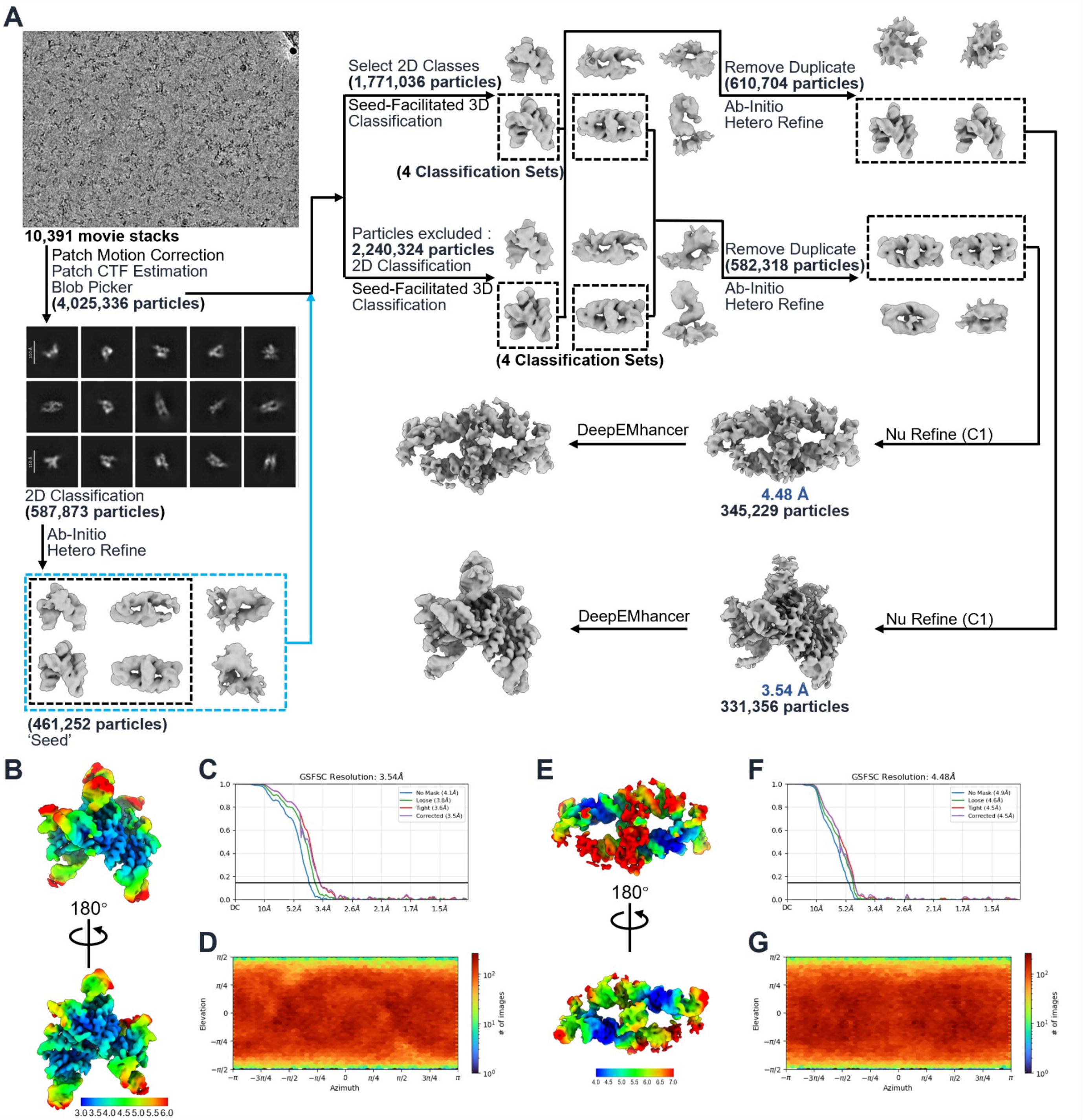


**Figure S21. Cryo-EM data processing workflow for EGFOA_*Pae*_6186.** (**A**) Flowchart of the single-particle cryo-EM data processing. (**B-G**) Local resolution maps (**B**, **E**), gold-standard FSC curves (**C**, **F**), and Euler angle distributions (**D**, **G**).

**Figure S22**

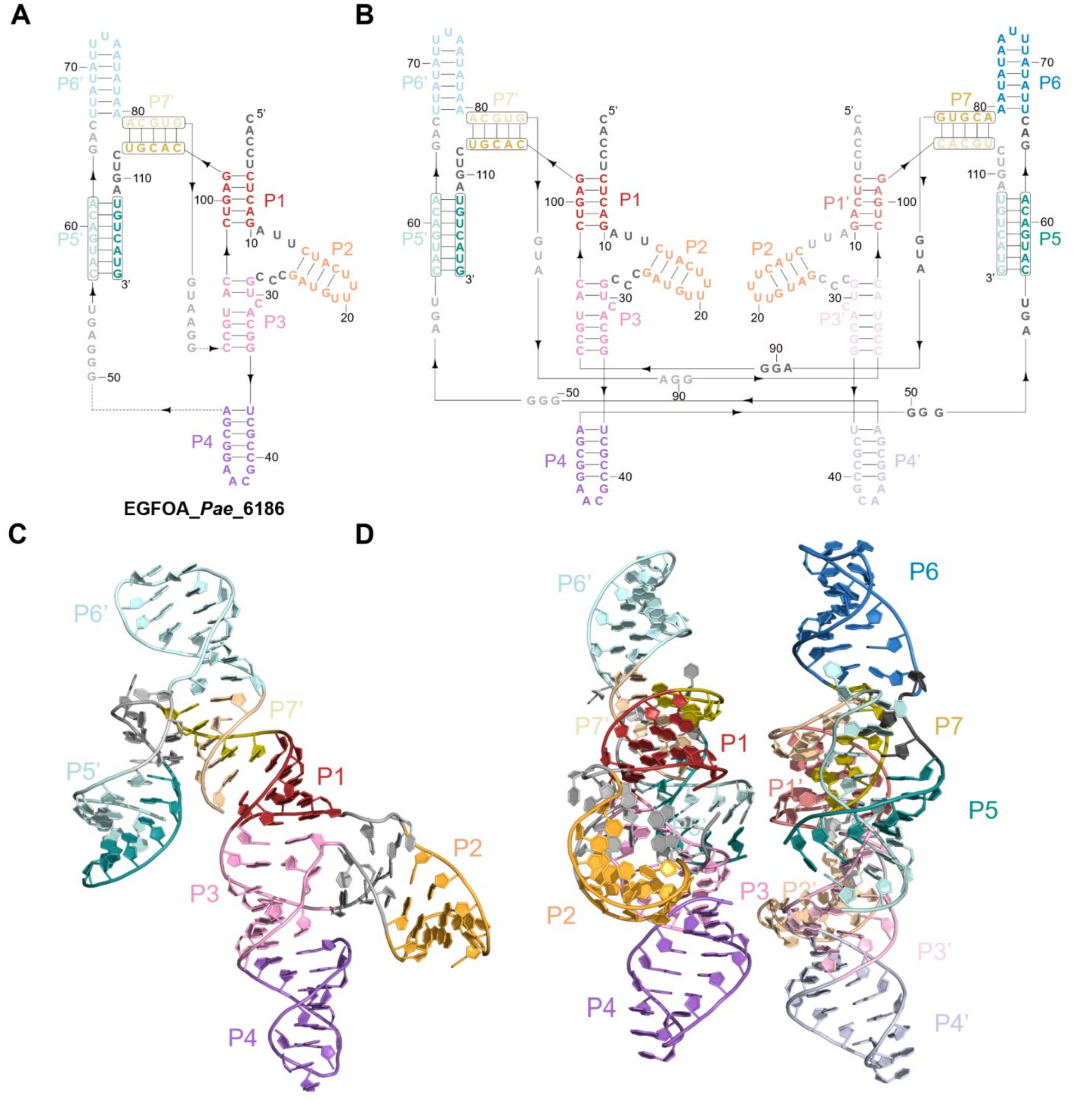


**Figure S22. Structural features and strand-displacement-mediated assembly of EGFOA_*Pae*_6186 RNA.** (**A**) Secondary structure of the EGFOA_*Pae*_6186 monomeric functional unit, with paired regions and loops distinctively colored. The lighter-colored sequence at the 3′ end represents a separate RNA chain, indicating that the functional unit incorporates a segment from an adjacent molecule. (**B**) Secondary structure diagram of the EGFOA_*Pae*_6186 dimer, illustrating the intermolecular 3′-end strand displacement mechanism that drives dimerization. (**C**) 3D atomic model of the monomeric functional unit, color-coded to match (**A**). (**D**) Overall 3D structure of the EGFOA_*Pae*_6186 dimer. The constituent interacting units are colored to correspond with the schematic in (**B**), visually highlighting the strand-swapped assembly interface.

**Figure S23**

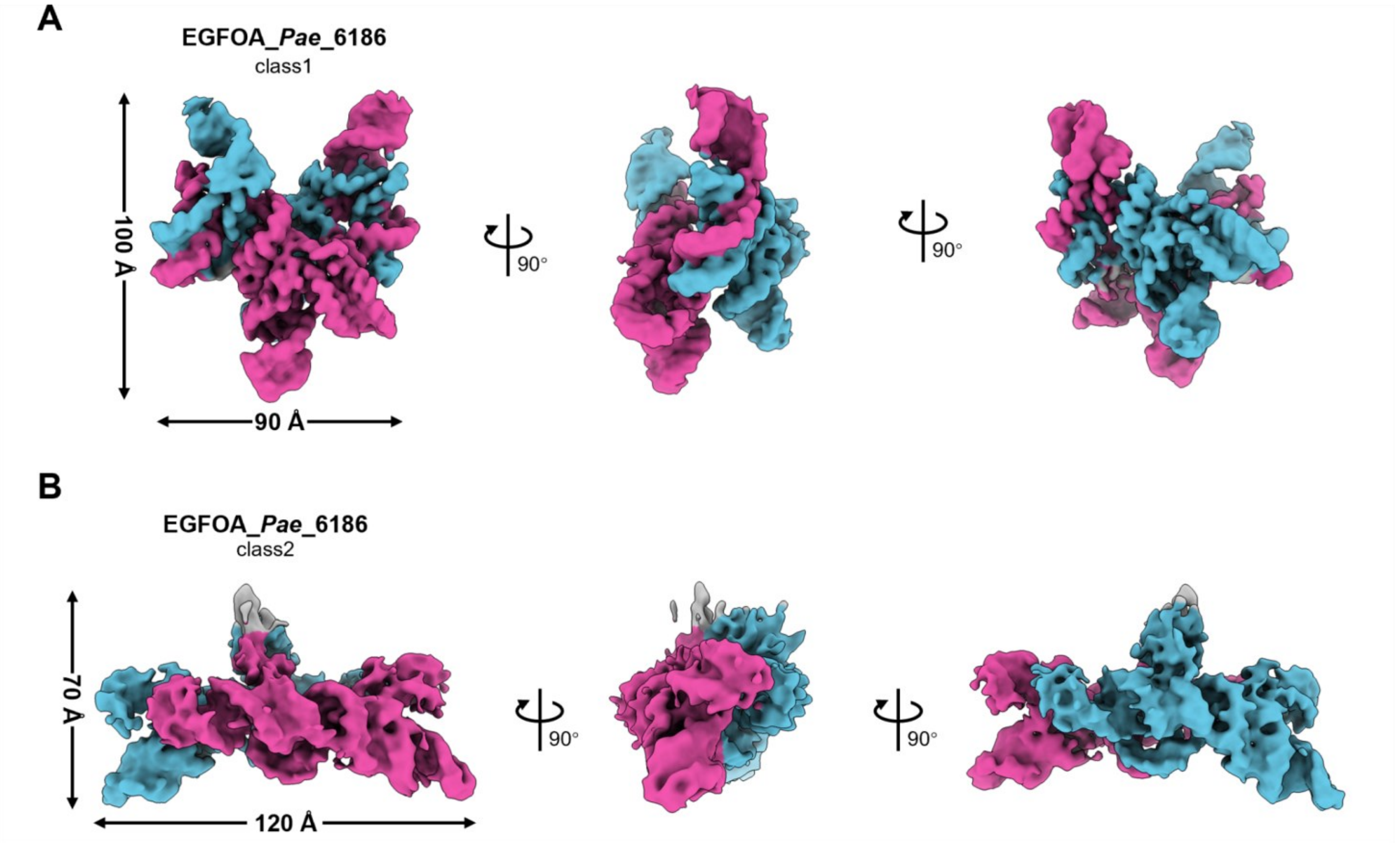


**Figure S23. Conformational flexibility and dynamic states of the EGFOA_*Pae*_6186 dimer.** Cryo-EM density maps illustrate two distinct structural conformations of the assembled dimer, highlighting its dynamic nature. Each conformational state is shown in three views (rotated by 90°), with the two interacting functional units distinctively colored in cyan and magenta. (**A**) The compact dimeric conformation (class 1) has overall dimensions of ~100 Å × 90 Å. (**B**) The extended dimeric conformation (class 2) demonstrates a significant structural rearrangement with overall dimensions of ~70 Å × 120 Å.

#### Figure S24

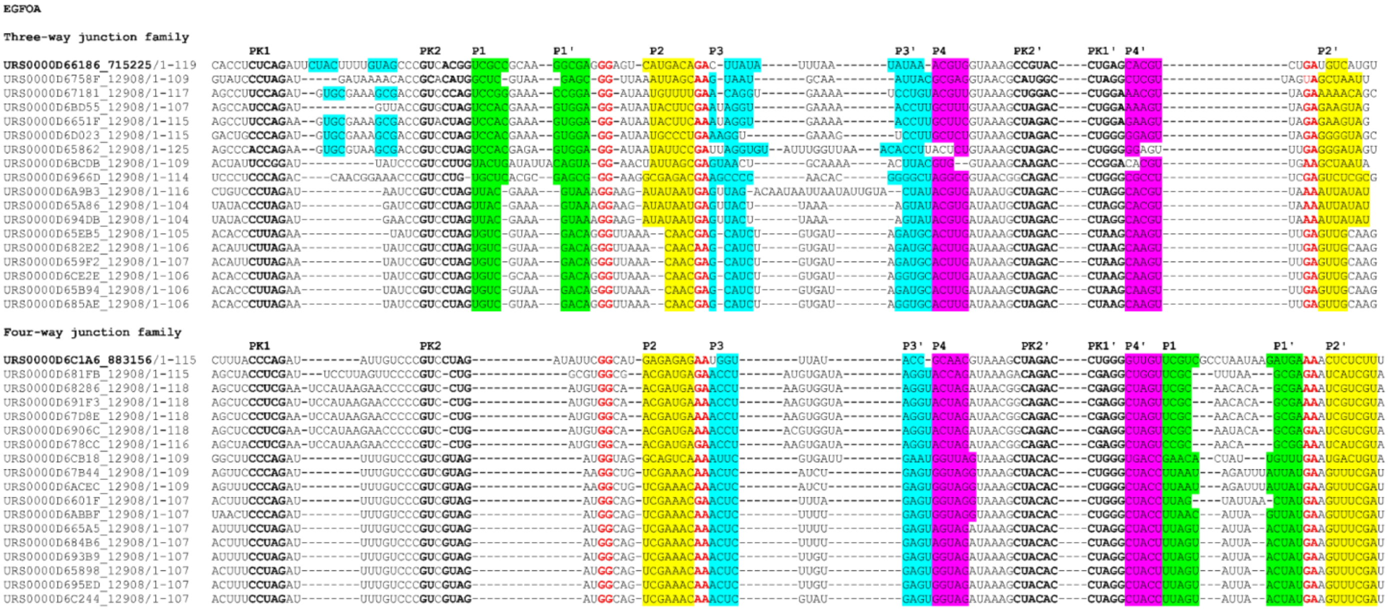

**Figure S24. Structure-based sequence alignment of the EGFOA RNA family.** Multiple sequence alignment of representative EGFOA RNA variants, illustrating sequence and structural conservation. On the basis of their secondary structure topologies, the family is phylogenetically classified into two major subfamilies: the three-way junction family (top) and the four-way junction family (bottom). Conserved structural elements, including canonical paired helices (P1–P4) and pseudoknots (PK1, PK2), are highlighted with distinct color-coded blocks. Highly conserved individual nucleotides are denoted in red text.

**Figure S25**

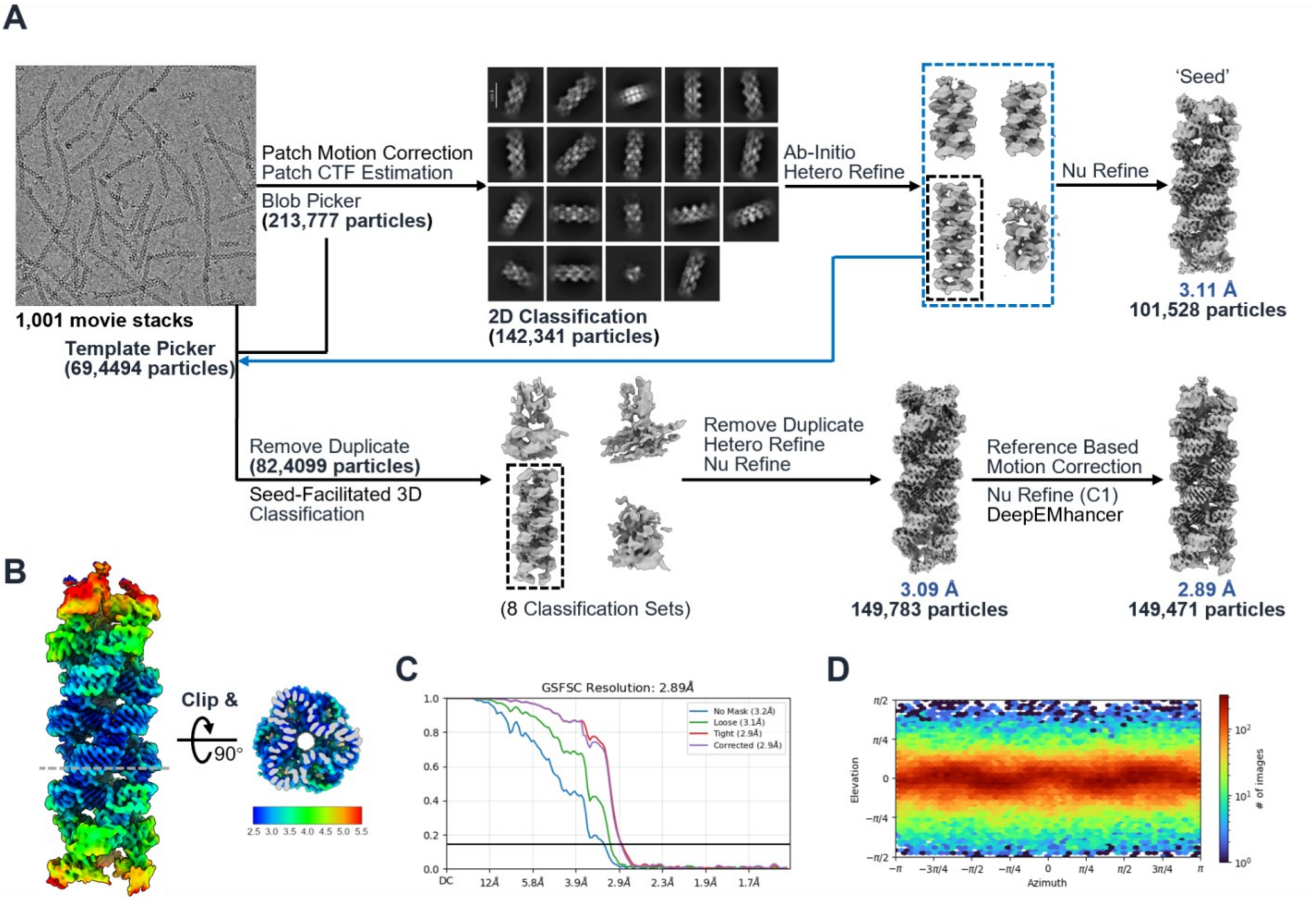


**Figure S25. Cryo-EM data processing workflow for the *Hm* kt7_6U RNA filament.** (**A**) Flowchart of the single-particle cryo-EM data processing. (**B**-**D**) Local resolution map (**B**), gold-standard FSC curves (**C**), and Euler angle distribution (**D**).

**Figure S26**

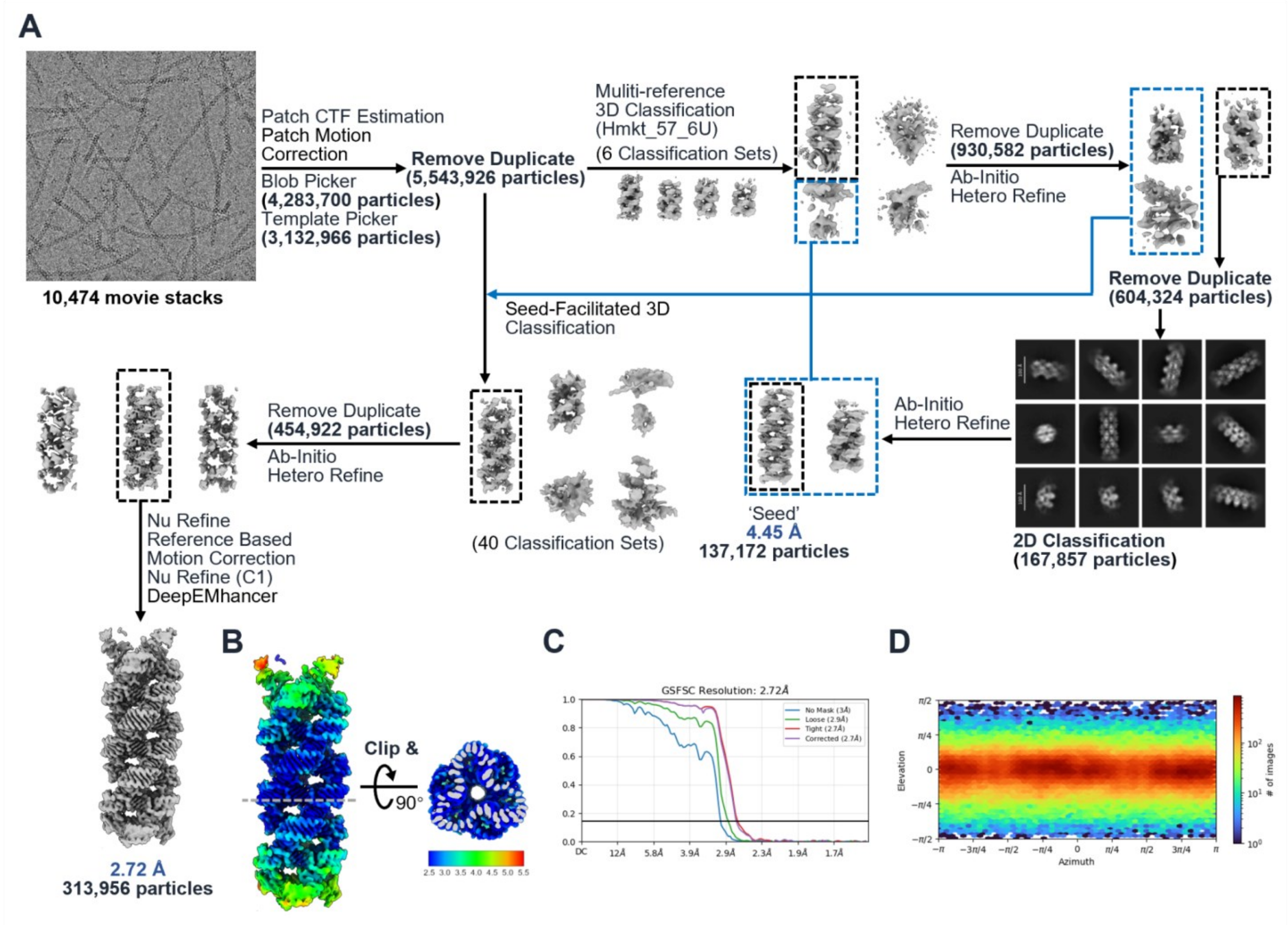


**Figure S26. Cryo-EM data processing workflow for the *Hm* kt7_5U RNA filament.** (**A**) Flowchart of the single-particle cryo-EM data processing. (**B**-**D**) Local resolution map (**B**), gold-standard FSC curves (**C**), and Euler angle distribution (**D**).

**Figure S27**

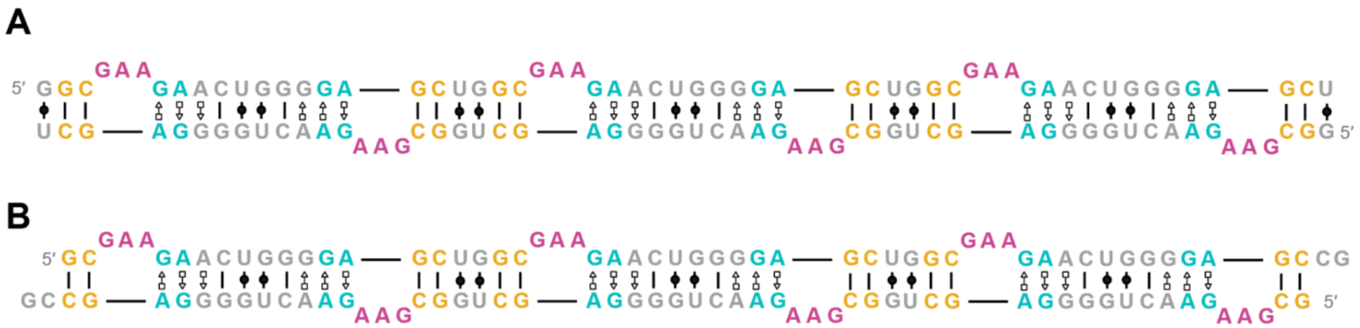


**Figure S27. Sequence and secondary structure schematics of the *Hm* kt7-57 RNA constructs.** (**A**, **B**) Detailed 2D sequence and secondary structure diagrams for the *Hm* kt7-57-6U (**A**) and *Hm* kt7-57-5U (**B**) RNAs used for the filament assemblies.

**Figure S28**

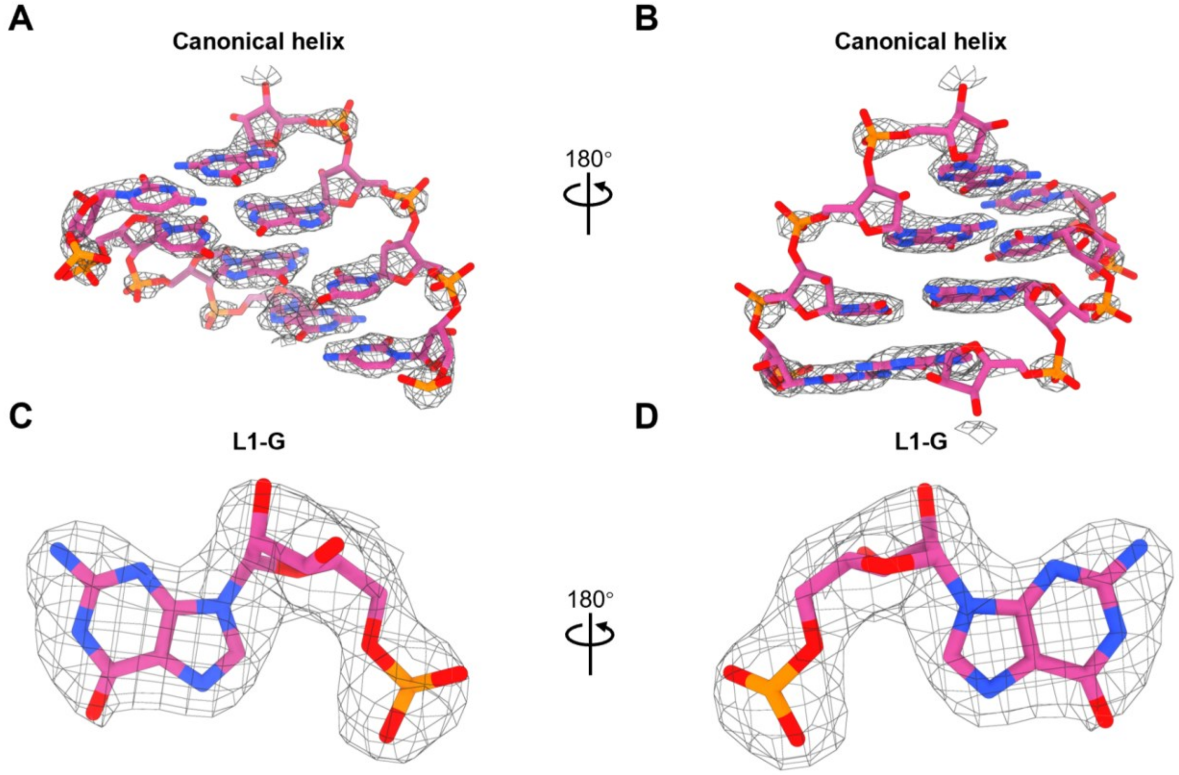


**Figure S28. High-resolution cryo-EM density map of the *Hm* kt7-5U RNA filament at 2.72 Å.** Representative cryo-EM density maps (displayed as gray mesh) are superimposed with the refined atomic model (shown in stick representation). (**A**, **B**) Two opposite views (rotated by 180°) of the canonical helical. (**C**, **D**) Two opposite views (rotated by 180°) of a single representative nucleotide (L1-G).

**Figure S29**

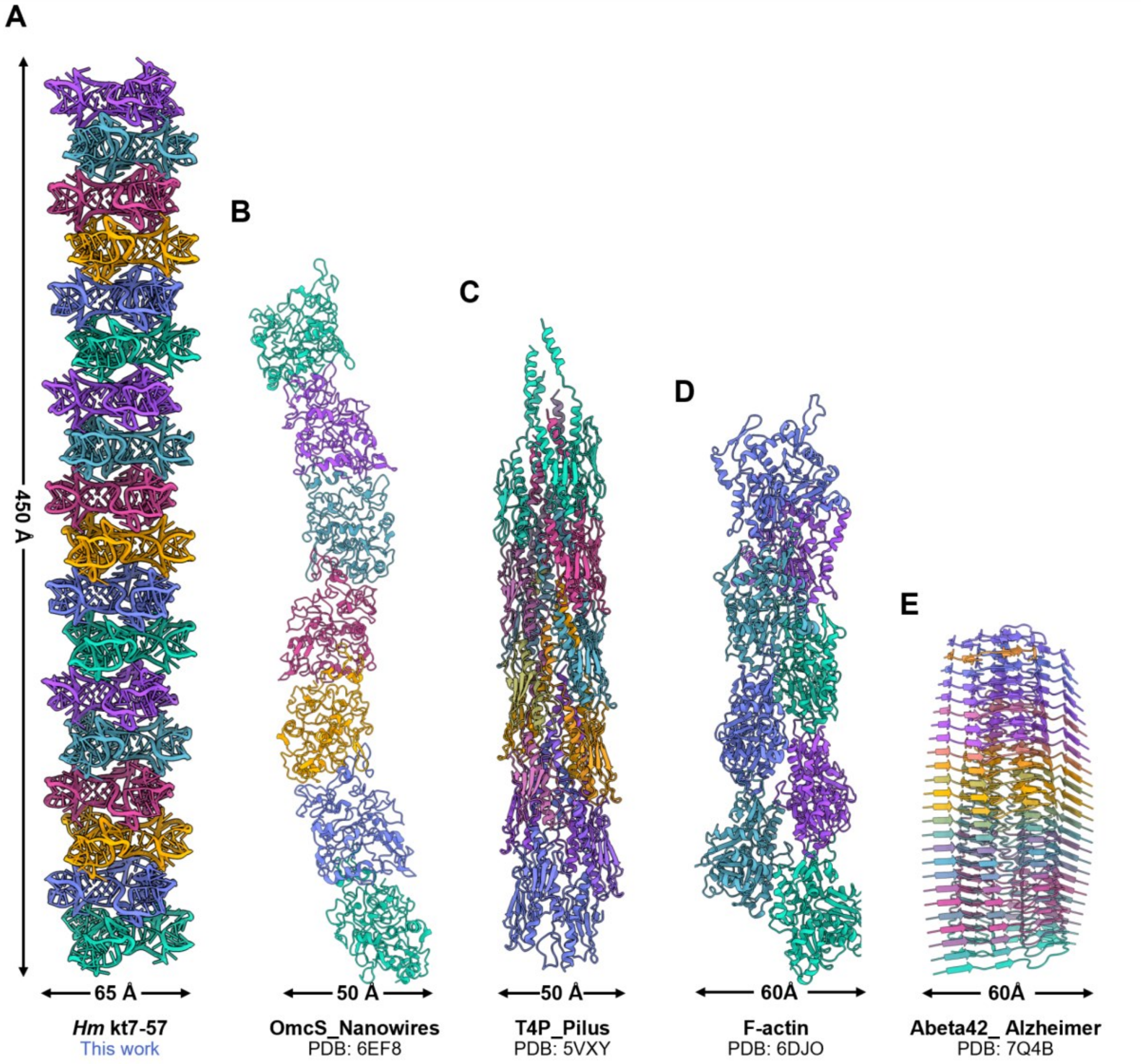


**Figure S29. Structural comparison of the *Hm* kt7-57 RNA filament with biological protein filaments.** (**A**) Longitudinal side view of the *Hm* kt7-57 RNA filamentous assembly (this work). (**B**–**D**) Side views of representative naturally occurring protein filaments, including (**B**) the electrically conductive OmcS nanowire (PDB: 6EF8)[4], (**C**) the Type IV pilus (T4P; PDB: 5VXY)[5], (**D**) the cytoskeletal F-actin filament (PDB: 6DJO)[6] and (**E**) the pathogenic amyloid-beta 42 (Aβ42) fibril associated with Alzheimer's disease (PDB: 7Q4B)[7]. All structures are displayed in cartoon representation, with individual constituent monomers or repeating asymmetric units distinctively colored to illustrate the helical or stacked structures.

**Figure S30**

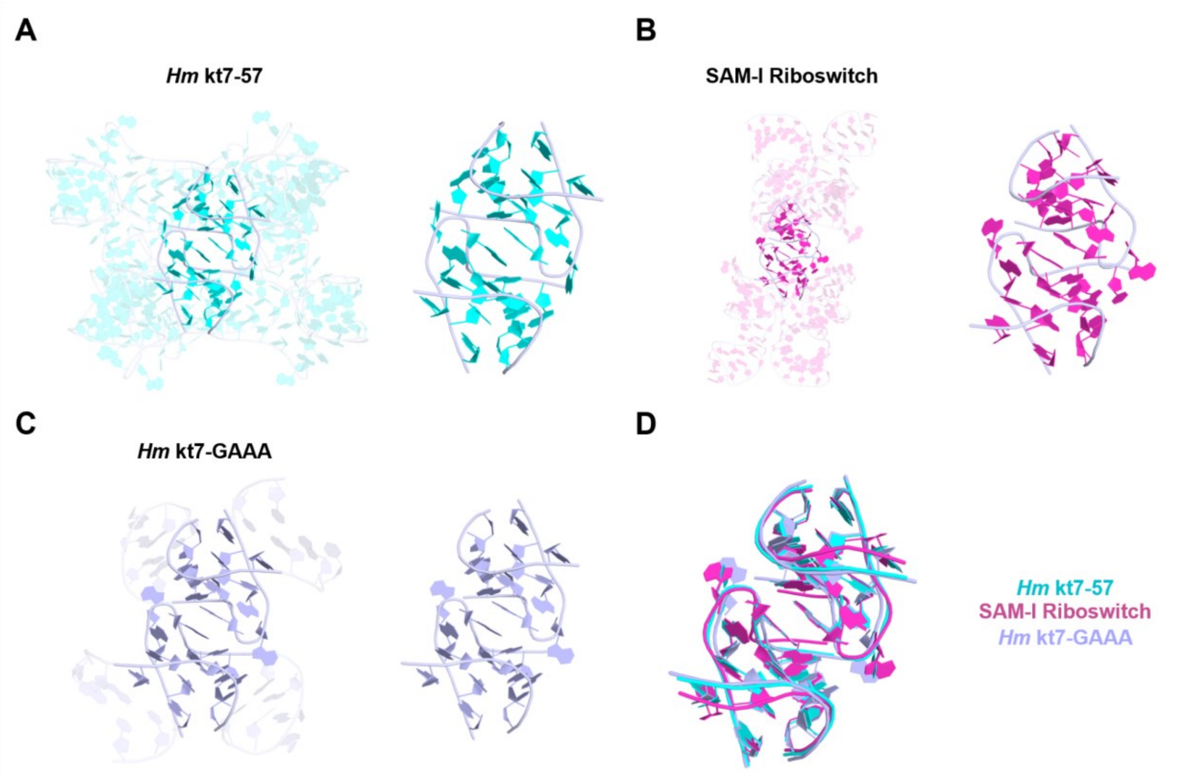


**Figure S30. Structural conservation of the kink-turn interaction motif across diverse RNAs.** (**A-D**) Overall 3D architectures and isolated views of the core kink-turn (k-turn) interaction modules. The broader RNA contexts are rendered transparently to highlight the specific k-turn motifs (shown as solid ribbons and bases) for *Hm* kt7-57 (cyan) (**A**), the crystal structure of the SAM-I riboswitch (magenta) (**B**), and the crystal structure of *Hm* kt7-GAAA (light blue) (**C**). (**D**) Structural superposition of the isolated k-turn motifs from the three distinct RNAs, demonstrating the high conformational conservation and structural rigidity of this interaction interface across different functional contexts.

**Figure S31**

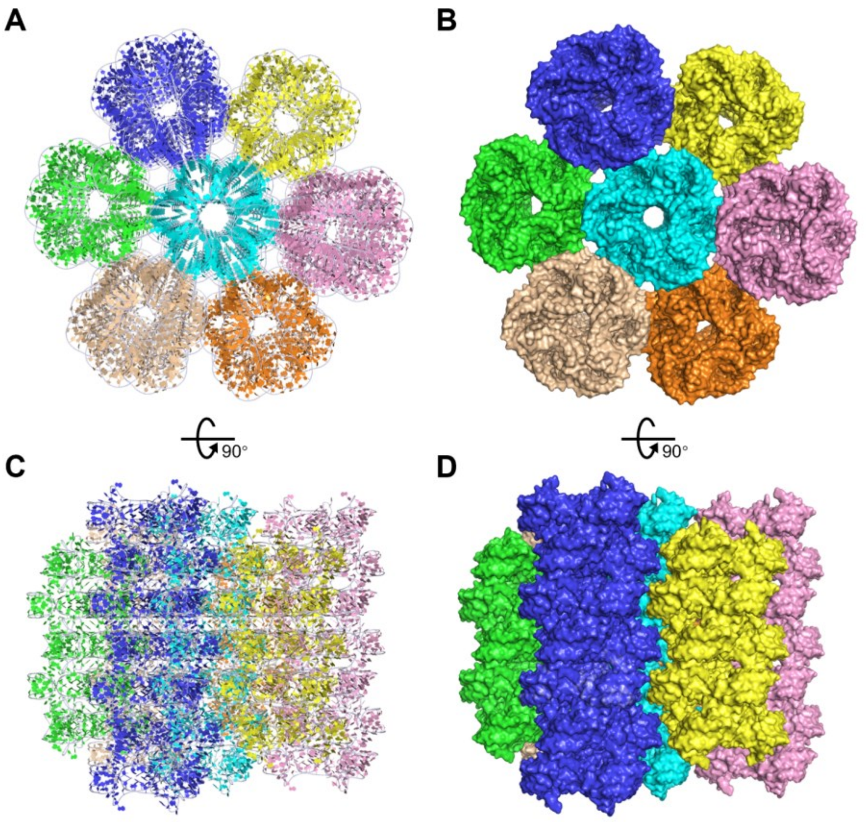


**Figure S31. Crystal packing and higher-order bundling of the *Hm* kt7-57-6U RNA (PDB: 5G4T).** (**A**) Axial view of the crystal packing lattice. (**B**) Surface representation of the axial view shown in (**A**). (**C**) Longitudinal side view (rotated by 90° from the axial view) of the crystal packing. (**D**) Surface representation of the longitudinal side view shown in (**C**).

**Figure S32**

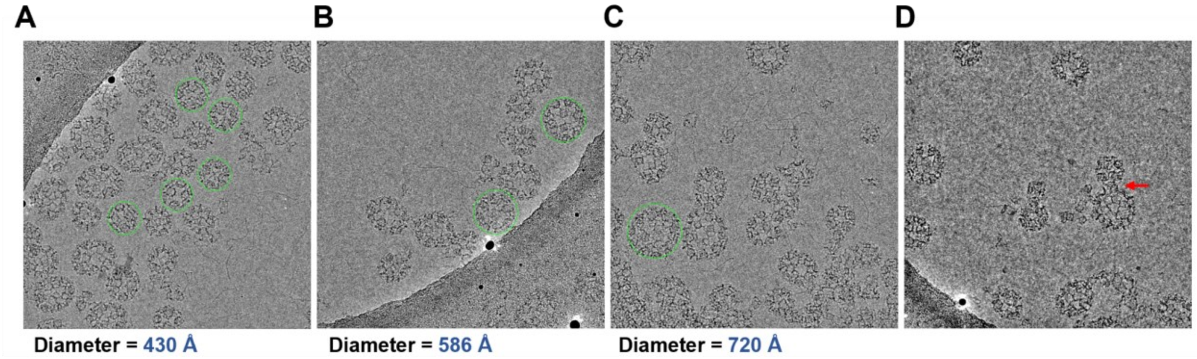


**Figure S32. Morphological heterogeneity and expanded assembly capacity of the manA_A0F1 RNA. (A)** Representative cryo-EM micrograph highlighting the predominantly observed spherical particles (green circles) with a diameter of ~430 Å, which correspond to the reconstructed 60-subunit icosahedral assembly. (**B**, **C**) Micrographs from the same dataset revealing the formation of even larger, higher-order spherical architectures, with representative particle diameters of approximately 586 Å (**B**) and 720 Å (**C**) indicated by green circles. (**D**) A representative field of view capturing a potential intermediate structural state (indicated by a red arrow), suggestive of dynamic fusion or fission events between the RNA assemblies.

**Figure S32**

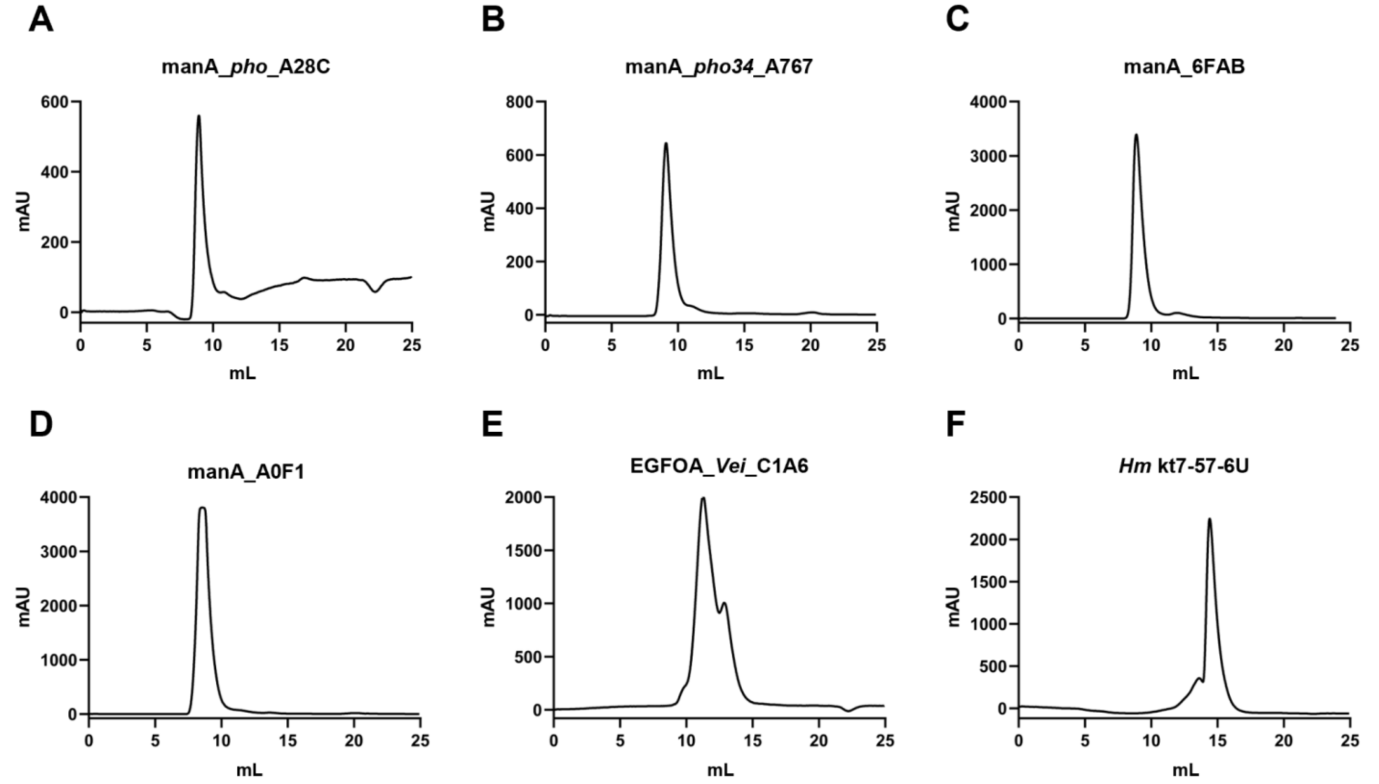


**Figure S32. Size-exclusion chromatography (SEC) purification profiles of the assembled RNA constructs.** UV absorbance traces (mAU) are plotted against the elution volume (mL) to assess the homogeneity of the assembled RNAs prior to cryo-EM grid preparation. (**a-d**) SEC chromatograms of the manA series RNA variants: (**A**) manA_*pho*_A28C, (**B**) manA_*pho34*_A767, (**C**) manA_6FAB, and (**D**) manA_A0F1. The sharp and symmetric elution peaks are indicative of highly uniform, discrete higher-order assemblies in solution. (**E**) SEC profile of the EGFOA_*Vei*_C1A6 RNA. The broader elution peak with visible shoulders reflects the coexistence of multiple oligomeric states (trimer, tetramer, and hexamer), perfectly consistent with the dynamic structural polymorphism observed in the cryo-EM data. (**F**) SEC chromatogram of the *Hm* kt7-57-6U RNA filament.

#### Table S1

| | manA_*pho*_A28C decamer (EMDB-69109) (PDB 23NP) | manA_*pho*_A28C pentamer (EMDB-69130) (PDB 23OO) | manA_*pho34*_A767 decamer (EMDB-69111) (PDB 23NR) | manA_*pho34*_A767 pentamer (EMDB-69112) (PDB 23NS) |
|---|---|---|---|---|
| **Data collection and processing** | | | | |
| **Microscope** | Titan Krios | | | |
| **Camera** | Falcon 4i | | Falcon 4i | |
| **Spherical Aberration (mm)** | 0.001 | | 2.7 | |
| Magnification | 165,000 | | 215,000 | |
| Voltage (kV) | 300 | | 300 | |
| Electron exposure (e–/Å$^2$) | 50 | | 50 | |
| Defocus range (µm) | -0.8 ~ -1.6 | | -0.8 ~ -1.6 | |
| Pixel size (Å) | 0.584 | | 0.572 | |
| Symmetry imposed | C5 | D5 | C5 | D5 |
| Initial particle images (no.) | 3,378,396 | | 1,730,049 | |
| Final particle images (no.) | 401,234 | 13,965 | 130,429 | 11,011 |
| Map resolution (Å) | 2.8 | 4.5 | 2.7 | 3.3 |
| FSC threshold | 0.143 | 0.143 | 0.143 | 0.143 |
| **Refinement** | | | | |
| Initial model used (PDB code) | De novo | manA_pho_A28C-C5 | De novo | manA_pho34_A767-C5 |
| Model resolution (Å) | 2.9 | 4.4 | 2.9 | 3.3 |
| FSC threshold | 0.5 | 0.5 | 0.5 | 0.5 |
| Map sharpening *B* factor (Å$^2$) | -74.4 | -130.7 | -59.0 | -53.8 |
| Model composition | | | | |
| Non-hydrogen atoms | 16,540 | 33,080 | 16,545 | 33,090 |
| RNA residues | 775 | 1,550 | 775 | 1,550 |
| Ligands | N/A | N/A | N/A | N/A |
| *B* factors (Å$^2$) | | | | |
| RNA | 50.07 | 37.80 | 58.50 | 83.10 |
| Ligand | N/A | N/A | N/A | N/A |
| R.m.s. deviations | | | | |
| Bond lengths (Å) | 0.007 | 0.007 | 0.006 | 0.008 |
| Bond angles (°) | 0.731 | 0.847 | 0.708 | 0.717 |
| Validation | | | | |
| MolProbity score | 2.70 | 3.25 | 2.72 | 2.77 |
| Clashscore | 10.77 | 41.70 | 11.25 | 12.84 |
| Poor rotamers (%) | 0 | 0 | 0 | 0 |
| $C_{mask}$ | 0.80 | 0.70 | 0.80 | 0.80 |

**Table S1(Continued).**

| | manA_A0F1 monomer (EMDB-69755) (PDB 24QD) | manA_A0F1 composite map (EMDB-69756) (PDB 24QE) | manA_67AA monomer (EMDB-69113) (PDB 23NT) | manA_6FAB monomer (EMDB-69108) (PDB 23NO) |
|---|---|---|---|---|
| **Data collection and processing** | | | | |
| **Microscope** | Titan Krios | | | |
| **Camera** | Falcon 4i | | Falcon 4i | Gatan K3 Summit |
| **Spherical Aberration (mm)** | 2.7 | | 2.7 | 2.7 |
| Magnification | 130,000 | | 215,000 | 215,000 |
| Voltage (kV) | 300 | | 300 | 300 |
| Electron exposure (e–/Å$^2$) | 50 | | 50 | 50 |
| Defocus range (μm) | -0.8 ~ -1.6 | | -0.8 ~ -1.6 | -0.8 ~ -1.6 |
| Pixel size (Å) | 0.934 | | 0.572 | 0.646 |
| Symmetry imposed | C1 | I | C1 | C1 |
| Initial particle images (no.) | 526,948 | | 1,510,114 | 3,176,047 |
| Final particle images (no.) | 420,841 | 420,841 | 384,770 | 761,723 |
| Map resolution (Å) | 3.3 | 3.3* | 3.4 | 3.0 |
| FSC threshold | 0.143 | 0.143 | 0.143 | 0.143 |
| **Refinement** | | | | |
| Initial model used (PDB code) | De novo | manA_A0F1-C1 | De novo | De novo |
| Model resolution (Å) | 3.5 | 3.7 | 3.2 | 3.1 |
| FSC threshold | 0.5 | 0.5 | 0.5 | 0.5 |
| Map sharpening *B* factor (Å$^2$) | -84.8 | -84.8 | -86.2 | -54.6 |
| Model composition | | | | |
| Non-hydrogen atoms | 3,338 | 200,280 | 2,757 | 2,477 |
| RNA residues | 156 | 9,360 | 129 | 115 |
| Ligands | N/A | N/A | N/A | N/A |
| *B* factors (Å$^2$) | | | | |
| RNA | 138.35 | 115.81 | 77.91 | 69.94 |
| Ligand | N/A | N/A | N/A | N/A |
| R.m.s. deviations | | | | |
| Bond lengths (Å) | 0.007 | 0.008 | 0.007 | 0.007 |
| Bond angles (°) | 1.029 | 1.045 | 1.111 | 1.344 |
| Validation | | | | |
| MolProbity score | 2.94 | 3.16 | 2.85 | 2.85 |
| Clashscore | 19.71 | 33.36 | 15.91 | 15.82 |
| Poor rotamers (%) | 0 | 0 | 0 | 0 |
| $CC_{mask}$ | 0.76 | 0.84 | 0.69 | 0.73 |

**Table S1(Continued).**

| | *Hm* kt7-57-6U filament (EMDB-69115) (PDB 23NU) | *Hm* kt7-57-5U filament (EMDB-69116) (PDB 23NV) | EGFOA_*Vei*_C1A6 trimer (EMDB-69107) (PDB 23NN) | EGFOA_*Pae*_6186 dimer (EMDB-69102) (PDB 23ND) |
|---|---|---|---|---|
| **Data collection and processing** | | | | |
| **Microscope** | Titan Krios | | | |
| **Camera** | Falcon 4i | Falcon 4i | Falcon 4i | Gatan K3 Summit |
| **Spherical Aberration (mm)** | 2.7 | 2.7 | 2.7 | 2.7 |
| Magnification | 165,000 | 165,000 | 215,000 | 215,000 |
| Voltage (kV) | 300 | 300 | 300 | 300 |
| Electron exposure (e–/Å$^2$) | 50 | 50 | 50 | 50 |
| Defocus range (µm) | -0.8 ~ -1.6 | -0.8 ~ -1.6 | -0.8 ~ -1.6 | -0.8 ~ -1.6 |
| Pixel size (Å) | 0.731 | 0.731 | 0.572 | 0.646 |
| Symmetry imposed | C1 | C1 | C3 | C1 |
| Initial particle images (no.) | 824,099 | 5,543,926 | 1,851,159 | 4,025,336 |
| Final particle images (no.) | 149,471 | 313,956 | 107,800 | 331,356 |
| Map resolution (Å) | 2.9 | 2.7 | 3.0 | 3.5 |
| FSC threshold | 0.143 | 0.143 | 0.143 | 0.143 |
| **Refinement** | | | | |
| Initial model used (PDB code) | 5G4T | 5G4T | De novo | De novo |
| Model resolution (Å) | 3.0 | 2.9 | 3.1 | 3.9 |
| FSC threshold | 0.5 | 0.5 | 0.5 | 0.5 |
| Map sharpening *B* factor (Å$^2$) | -67.2 | -73.9 | -78.3 | -171.6 |
| Model composition | | | | |
| Non-hydrogen atoms | 15,012 | 15,012 | 6,129 | 4,327 |
| RNA residues | 684 | 684 | 288 | 204 |
| Ligands | N/A | N/A | N/A | N/A |
| *B* factors (Å$^2$) | | | | |
| RNA | 61.40 | 50.87 | 78.69 | 75.31 |
| Ligand | N/A | N/A | N/A | N/A |
| R.m.s. deviations | | | | |
| Bond lengths (Å) | 0.009 | 0.007 | 0.006 | 0.007 |
| Bond angles (°) | 0.689 | 0.822 | 1.256 | 1.770 |
| Validation | | | | |
| MolProbity score | 2.46 | 2.75 | 2.61 | 2.95 |
| Clashscore | 5.74 | 12.18 | 8.45 | 20.09 |
| Poor rotamers (%) | 0 | 0 | 0 | 0 |
| $CC_{mask}$ | 0.82 | 0.79 | 0.72 | 0.72 |

**Table S1. Cryo-EM data collection, refinement and validation statistics.**

*Reported resolutions are derived from composite maps

**Table S2**

| | Name | RNAcentral ID | Species | Resolution (Å) |
|---|---|---|---|---|
| **manA** | 6FAB | URS0000D66FAB_12908 | Unclassified | 3.00 |
| | A0F1 | URS0000D6A0F1_12908 | Unclassified | 3.40 |
| | 67AA | URS0000D667AA_12908 | Unclassified | 3.00 |
| | *pho*_A28C | URS0000D6A28C_314292 | *Photobacterium angustum* S14 | 2.80 |
| | *pho34*_A767 | URS0000D6A767_121723 | *Photobacterium* sp. SKA34 | 2.70 |
| | 6051 | URS0000D66051_12908 | Unclassified | ND |
| | 9027 | URS0000D69027_12908 | Unclassified | ND |
| | BB58 | URS0000D6BB58_12908 | Unclassified | ND |
| | 6849 | URS0000D66849_12908 | Unclassified | ND |
| | B1E1 | URS0000D6B1E1_12908 | Unclassified | ND |
| **EGFOA** | *Vei*_C1A6 | URS0000D6C1A6_883156 | *Veillonella seminalis* ACS-216-V-Col6b | 2.96 |
| | *Pae*_6186 | URS0000D66186_715225 | *Paenibacillus vortex* V453 | 3.54 |
| | *Pae*_D1F6 | URS0000D6D1F6_1087481 | *Paenibacillus peoriae* KCTC 3763 | ND |
| | D023 | URS0000D6D023_12908 | Unclassified | ND |
| | *Bac*_859C | URS0000D6859C_649639 | *Bacillus cellulosilyticus* DSM 2522 | ND |
| | B17B | URS0000D6B17B_12908 | Unclassified | ND |
| **RF02920** | 6FEE | URS0000D66FEE_12908 | Unclassified | ND |
| | D21C | URS0000D6D21C_12908 | Unclassified | ND |
| | C3EC | URS0000D6C3EC_12908 | Unclassified | ND |
| | 9644 | URS0000D69744_12908 | Unclassified | ND |
| **wcaG** | 7E1D | URS0000D67E1D_12908 | Unclassified | ND |
| | *Syn*_7E1D | URS0000D69BDD_238854 | *Synechococcus* phage S-PM2 | ND |
| | *Syn*_7DC0 | URS0000D67DC0_382359 | *Synechococcus* phage syn9 | ND |
| **FuFi-1** | A73B | URS0000D6A73B_12908 | Unclassified | ND |
| | *Fus*_A46B | URS0000D6A46B_469621 | *Fusobacterium* sp. 1_1_41FAA | ND |
| | *Clo*_87A6 | URS0000D687A6_999410 | *Clostridioforme* CM201 | ND |
| **RF03046** | *Aci*_71BD | URS0000D671BD_1221247 | *Acinetobacter baumannii* AB_1594-8 | ND |
| | *Aci*_9D41 | URS0000D69D41_575588 | *Acinetobacter lwoffii* SH145 | ND |
| | *Mor*_6CA6 | URS0000D66CA6_857572 | *Moraxella catarrhalis* 101P30B1 | ND |
| **RF02932** | *Bur*_65B0 | URS0000D665B0_441160 | *Burkholderia pseudomallei* 14 | ND |
| | *Tay*_B025 | URS0000D6B025_1008459 | *Taylorella asinigenitalis* MCE3 | ND |
| | *Azo*_CFDA | URS0000D6CFDA_748247 | *Azoarcus* sp. KH32C | ND |
| **RT-5** | *Clo*_C9A1 | URS0000D6C9A1_720554 | *Clostridium clariflavum* DSM 19732 | ND |
| | C_8965 | URS0000D68965_12908 | Unclassified | ND |
| | C_7A18 | URS0000D67A18_12908 | Unclassified | ND |
| ***Hm* kt7-57** | 5U | N/A | *Haloarcula marismortui* | 2.70 |
| | 6U | N/A | *Haloarcula marismortui* | 2.90 |

**Table S2. Summary of RNA sequences screened for cryo-EM.** N/A: Not Applicable; ND: not determined; Unclassified: unclassified sequences. Construct names comprise species abbreviations (if classified) followed by the last four characters of their RNAcentral IDs.

**Table S3**

| Name | Sequence (5' to 3') |
|---|---|
| *Hm*Kt7-57-5U | GCGAAGAACUGGGGAGCUGGCGAAGAACUGGGGAGCUGGCGAAGAACUGGGGAGCCG |
| *Hm*K-57-6U | GGCGAAGAACUGGGGAGCUGGCGAAGAACUGGGGAGCUGGCGAAGAACUGGGGAGCU |
| EGFOA_*Vei*_C1A6 | GGCUUUACCCAGAUAUUGUCCCGUCCUAGAUAUUCGGCAUGAGAGAGAAUGGUUUAUACCGCAACGUAAAGCUAGACCUGGGGUUGUUCGUCGCCUAAUAAGAUGAAAACUCUCUUU |
| EGFOA_*Pae*_6186 | GGCACCUCUCAGAUUCUACUUUUGUAGCCCGUCACGGUCGCCGCAAGGCGAGGGAGUCAUGACAGACUUAUAUUUAAUAUAAACGUGGUAAAGCCGUACCUGAGCACGUCUGAUGUCAUGU |
| manA_6FAB | GGUAAGGAAGUAGCUCUUCCUUAUCGGGUAUGGCCGAAUAAGUCCGCCACGGACUAAGGCACAGGGCACGCAAACCAAGAACCGACUGGUGGAGGAAUGCGUGACUGGUAACGCGUCUACACUGUAGUGGCAUAACGCGUUUGGAGGUAUACAGCAAGGCCUCCUACCCACCUU |
| manA_A0F1 | GGUGUUGGACGCAACAUGGGAGUGACUGAAUAAACUUACUGGCAACUGCUGGUUAAGGUGAUGAGACACAGGUGGUGCUGCUACGAAAGUAGAACCGAUCAACCAAUCGGGUCUCAGGCAAUAACGUUUUUACUACUGUAGUAAUGCCCGUUAUUUGUUGGUACACAGGAAUCCAACCUCCCUCCUU |
| manA_67AA | GGCGUGCCUGAUCAGCACAUUGGGUGUGACUGAAUAAACUUACUGGCAUAUAGCUGGUUAAGGUGAUACAACACAGGUGGUGCUGCUCCGAAAGGAGAAUCGACUUACCAGUCGGGUUGUAGGCAGAGAUGAUUUCUAAACUGUAGAAAUGCCCAUCUCUUGUUGGUAUACAGGAUUCCAACCACCCUCCCU |
| manA_*pho*_A28C | GGCUUUAGGCGACGGCUUGAAGCGGGGAGUGCAGAGAAUAAUAGUAUUCAGCGACUAUAAUCUGGCUUAUCACGGGUUGUGCCGACCUAACUUGCGGUGACGUUAAGUUAGGACAUCGCCGACAUGGCGGGAUUAAGGUUCAAUGAGAUCGGAGCUGACUCAUUGUUGUAGGUAUUCCAAAGUCCUACCUCCCACCUU |
| manA_*pho34*_A767 | GGCUUUAGACGACGGUUUGAAGCGGGGAGUGCAGAGAAUAAUAGUAAUCAGCGACUAUAAUCUGGCUUAUCACGGGUUGUGCCGACCUAACUUGCGGUGACGUUAAGUUAGGACAUCGCCGACAUGGCGGGAUUAAGGUUCGAUGAGAUCGGAGCUGACUCAUUGUUGUAGGUAUUCCAAAGUCCUACCUCCCACCUU |

**Table S3. RNA sequences utilized for structural determination.**
